# The mammalian genome is punctuated by regularly spaced nucleosome islands

**DOI:** 10.64898/2026.09.30.755582

**Authors:** M. Christophe, L. Altamirano, P. Navarro, J. Mozziconacci

## Abstract

Nucleosome positioning plays a fundamental role in regulating DNA accessibility and cellular activity, yet the precise rules by which mammalian sequences instruct this organization have remained incomplete. Prior work has established that intrinsic sequence preferences dictate nucleosome architecture in yeast, but in mammals, this sequence code is modulated by chromatin state and cell-type–specific factors. Here, we train a deep-learning model on genome-wide maps from mouse embryonic stem cells to predict nucleosome occupancy directly from the primary DNA sequence. Our model accurately predicts *in vivo* occupancy and captures shared biological signals while effectively denoising assay-specific technical biases. Through in silico mutagenesis, we identify specific nucleosome positioning regions comprised of short 5-to-20-base-pair motifs concentrated within dense regulatory windows. Furthermore, we demonstrate that repetitive DNA elements play a major role in establishing these highly organized nucleosome arrays on a genomic scale. Specifically, CCCTC-binding factor motifs embedded within short interspersed nuclear elements, zinc-finger–bound motifs within long interspersed nuclear elements, and microsatellite repeats act as potent organizers of nucleosome phasing. Together, our findings extend the concept of sequence-encoded nucleosome architecture to mammalian genomes and reveal that regularly spaced nucleosome islands are shaped by short, recurrent sets of DNA motifs that couple local sequence to chromatin structure.

## Introduction

Nucleosomes, the fundamental units of chromatin, package 147 bp of DNA around histone octamers. In addition to compacting the genome and contributing to its three-dimensional structure (Routh et al., 2008; Correll et al., 2012; Teif and Bohinc, 2011), nucleosomes are a conserved hallmark of genome organization and gene regulation, because their positions modulate DNA accessibility to Transcription Factors (TFs), polymerases and chromatin remodelers (Kornberg and Lorch, 2020). In multicellular organisms, nucleosome positions differ between cell types and along differentiation trajectories and genome-wide nucleosome maps provide a readout of cell-type–specific genome activity (Snyder et al., 2016). This raises a central question: to what extent does the DNA sequence itself instruct nucleosome organization in mammalian genomes, and can we exploit nucleosome maps to decode underlying regulatory information? Pioneering studies in budding yeast established that intrinsic DNA sequence preferences make a major contribution to nucleosome positioning *in vivo* (Segal et al., 2006; Kaplan et al., 2009). Comparing nucleosomes assembled *in vitro* on purified genomic DNA with high-resolution *in vivo* maps showed that dinucleotide periodicities and sequence composition over 5-bp words are sufficient to explain a large fraction of the yeast nucleosome landscape and to construct a predictive probabilistic model of nucleosome occupancy. Expanding these results, convolutional neural networks were recently trained to predict MNase-seq nucleosome density across the Saccharomyces cerevisiae genome and to compute a “mutation score” for every base, revealing that poly(A) and CG-rich motifs—often associated with the RSC remodeler—are key sequence features controlling nucleosome exclusion in Nucleosome Depleted Regions (NDRs) upstream of transcription start sites (Routhier et al., 2021). These yeast studies demonstrate that deep models can learn a compact representation of the “nucleosome code” directly from sequence and use it to interpret the impact of mutations on chromatin structure.

Genome-wide MNase-seq maps in mammals have revealed a more heterogeneous picture, in which the influence of DNA sequence is modulated by chromatin state and cell type. In primary human granulocytes and T cells only a minority of the genome harbors strongly positioned nucleosomes, whereas most regions show flexible occupancy, and identified GC-rich “container sites” whose sequence-encoded affinity nucleates phased nucleosome arrays with cell-type–specific linker lengths (Valouev et al., 2011). In human lymphoblastoid cells nearly half of the genome participates in regularly spaced nucleosome arrays, and these arrays arise from an interplay between sequence preferences and barriers created by DNA-binding proteins and other chromatin factors (Gaffney et al., 2012). Among these factors, the architectural protein CTCF (CCCTC-binding factor) acts as a primary physical barrier that establishes nucleosome phasing across the genome (Fu et al., 2008). During mouse embryonic stem cell differentiation nucleosome repositioning is tightly linked to DNA methylation thereby controlling a class of CTCF sites that are occupied in embryonic stem cells but lose CTCF in differentiated cells (Teif et al., 2014). CTCF further organizes local chromatin by imposing phased nucleosome arrays and locally reduced nucleosome repeat length in mouse embryonic stem cells (Clarkson et al., 2019; Owens et al., 2019). Other factors have also been identified as nucleosome phasing factors, both in interphase and mitosis (Festuccia et al., 2019). Together, these mammalian studies indicate that sequence-encoded nucleosome preferences (Gaffney et al., 2012), architectural proteins and epigenetic modifications jointly shape nucleosome organization in vivo, but they do not yet provide a comprehensive sequence-level description of the motifs that underlie strongly phased nucleosome patterns in vertebrate genomes.

Despite the availability of high-coverage MNase-seq and related chemical cleavage maps in mice and humans, most sequence-based analyses in mammals have relied on relatively simple models—such as k-mer enrichment, GC-content (Wang and Widom, 2005; Segal et al., 2006; Kaplan et al., 2009), or yeast-trained energetics models—to explain local nucleosome occupancy and spacing (Miele et al., 2008; Xi et al., 2010; Zhang et al., 2011; Chereji et al., 2011; Riposo and Mozziconacci, 2012), and have not systematically decomposed the contribution of short motifs to strongly phased nucleosome arrays. At the same time, yeast deep-learning models have so far been applied mainly within S. cerevisiae or in limited cross-species settings, and it remains unclear whether similar approaches can uncover distinct, mammal-specific sequence features that organize nucleosome arrays at regulatory elements and across repetitive elements (Di Gangi et al., 2018; Amato et al., 2020; Zhou et al., 2022; Han et al., 2022). Here we address these questions by training deep-learning models on genome-wide nucleosome maps in mouse embryonic stem cells, and by using interpretability analyses to extract the underlying DNA features that drive the model predictions. Building on the methodological framework developed in yeast (Routhier et al., 2021), we first show that a simple convolutional architecture can predict MNase-derived (Festuccia et al., 2019) and chemical cleavage–derived (Voong et al., 2016) nucleosome occupancy from the mouse genomic sequence and that the same sequence rules are sufficient to explain both assays. We then identify “phased nucleosome islands,” defined as regions containing on the order of ten well-positioned nucleosomes assembled around compact Nucleosome Positioning Regions (NPRs), which are themselves formed by 5–20 bp motifs concentrated within 100 bp windows. These NPRs are enriched at active transcription start sites and DNase I–accessible regions of the genome reflecting their potential functional role. Finally, we uncover a prominent association of repetitive DNA with these islands at a genome-wide scale, with major contributions from CTCF motifs embedded in Short Interspersed Nuclear Elements (SINEs), Zinc Finger (ZNF)-bound motifs within Long Interspersed Nuclear Elements (LINEs), and microsatellite repeats whose nucleosome-organizing effect depends on repeat copy number. Together, our results extend the concept of a sequence-encoded nucleosome organization from yeast to a mammalian genome and reveal that regularly spaced nucleosome islands are shaped by a small set of recurrent DNA motifs and repeat families that couple local sequence composition to chromatin structure in embryonic stem cells.

## Results

### The same sequence features explain both MNase-seq and chemical cleavage maps

To test whether nucleosome organization in mouse embryonic stem cellss (mESCs) can be inferred directly from the underlying DNA sequence, we trained a convolutional neural network similar in architecture to the deep-learning framework previously applied in S. cerevisiae for predicting MNase-seq nucleosome density and computing mutation-level effects Routhier et al. (2021). The model takes as input 2001-bp genomic windows and outputs a predicted nucleosome-density track (Fig. 1A, top). To interpret which sequence features drive these predictions, we performed *In Silico* Mutagenesis (ISM), in which each nucleotide is systematically substituted by the three alternative bases and the resulting change in predicted nucleosome density is aggregated into a per-position mutational sensitivity score (Fig. 1A, bottom).

**Figure 1.**
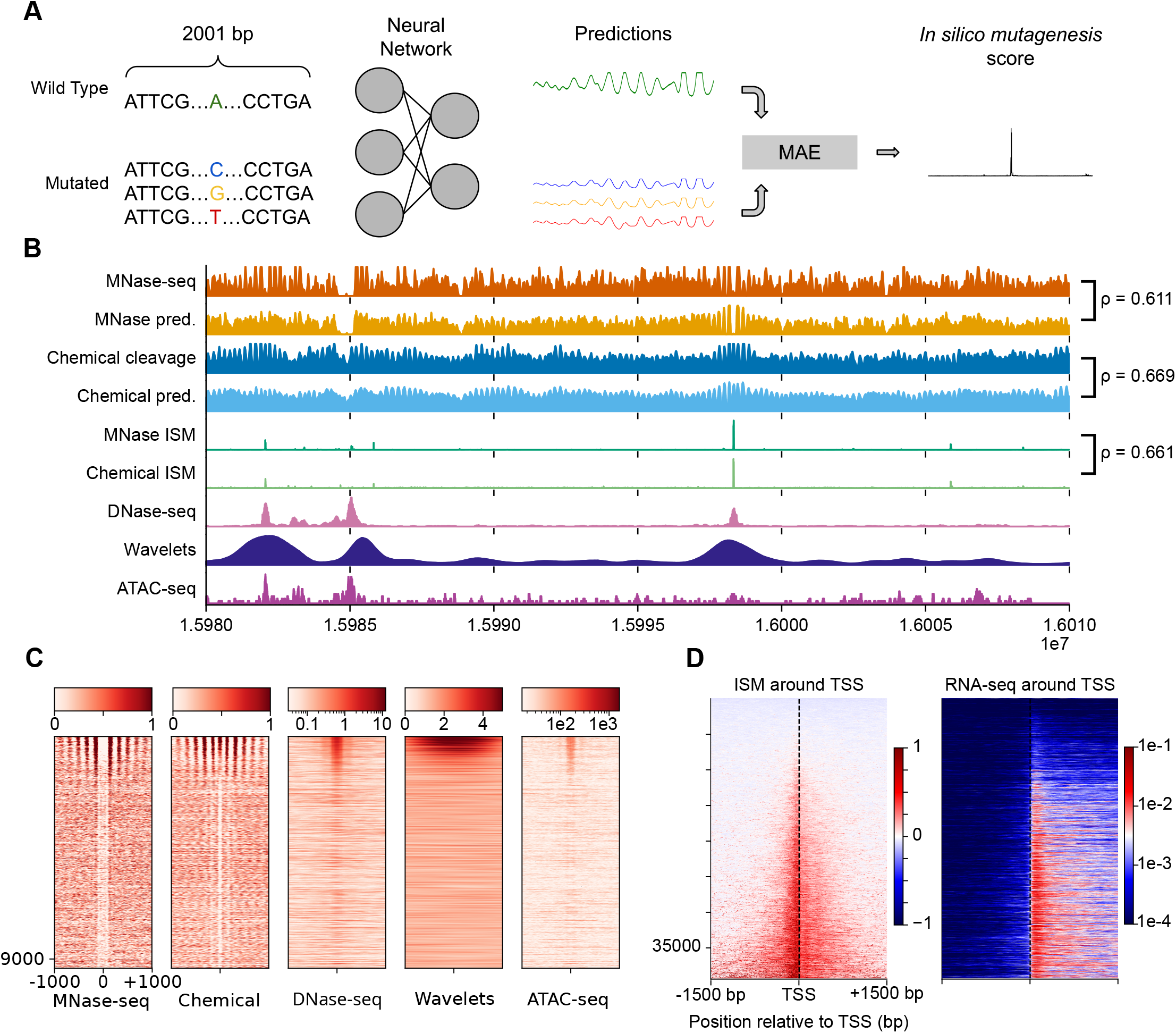
Interpretation of convolutional neural networks reveals nucleosome positioning regions that correlate with transcriptional and epigenetic features. (A) Schematic representation of the *in silico* mutagenesis (ISM) workflow. The neural network, trained on baseline nucleosome occupancy, predicts profiles for wild-type and mutated sequences; the resulting divergence is quantified by the Mean Absolute Error (*MAE*), defining the positional importance score. (B) Genomic locus (chr19:15,980,000-16,010,000) displaying predicted nucleosome occupancy tracks and orthogonal experimental signals. Genome-wide correlations are computed using Pearson’s correlation coefficient (*ρ*). (C) Heatmaps of Nucleosome occupancy, wavelets and chromatin accessibility signal intensities at identified ISM peaks. Each row represents a 2-kb genomic window centered on the ISM peak (*n* = 388, 694). (D) Stratification of ISM scores as a function of transcriptional activity. Intensities are plotted in the vicinity of promoters (3-kb window) and oriented according to the direction of transcription (*n* = 39385 promoters).

**Supplementary Figure 1.**
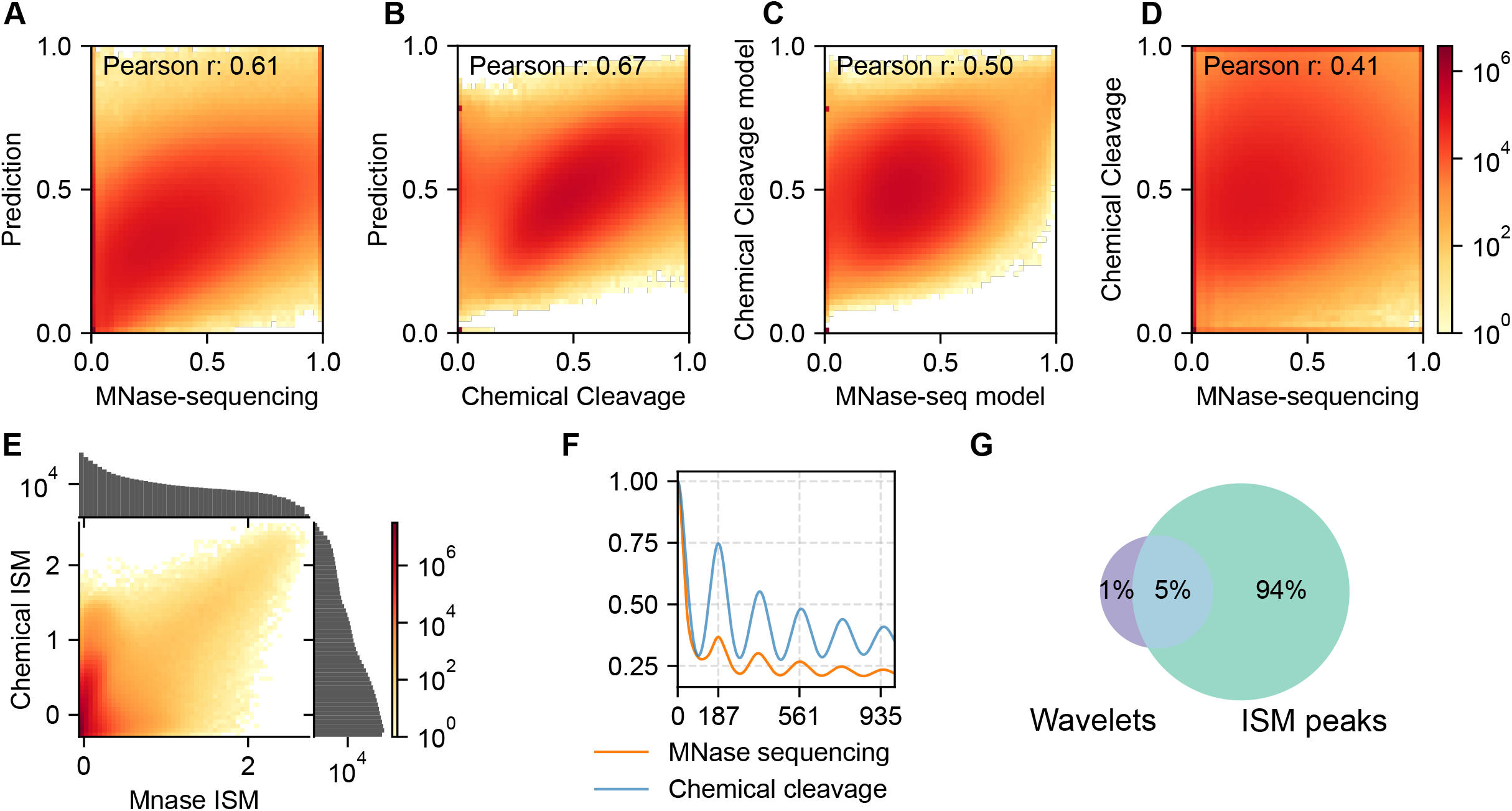
Cross-validation of deep learning models and generalization within low-mappability genomic regions. (**A–D**) Genome-wide scatter plots and Pearson correlation coefficients (*r*) comparing experimental versus predicted nucleosome occupancy profiles: (**A**) Experimental MNase-seq versus MNase-seq model predictions (*r* = 0.61).012 (B) Experimental chemical cleavage versus chemical cleavage model predictions (*r* = 0.67). (C) Predictions from the independently trained MNase-seq model versus the chemical cleavage model (*r* = 0.50). (**D**) Experimental MNase-seq tracks versus experimental chemical cleavage tracks (*r* = 0.41). (**E**) Joint distribution of MNase and Chemical ISM scores on chromosome 1, expressed as log_10_(z-score + 1) (*r* = 0.66). (**F**) Spatial autocorrelation profiles computed from experimental nucleosome occupancy data. (**G**) Venn diagram of the intersection between nucleosome island (blue) and ISM hotspots (green).

Using this sequence-based model, predicted nucleosome occupancy correlates strongly with experimental MNase-seq and chemical cleavage profiles (Fig. 1B). Model-to-experiment genomewide correlations reach 0.61 for MNase-seq and 0.67 for chemical cleavage (Fig. 1B, SUPP 1A,B), indicating that DNA-encoded rules underlie both assays. Notably, the predictions for MNase-seq and chemical cleavage correlate more strongly with each other (0.5, Fig.SUPP 1C) than the experimental datasets themselves (0.4, Fig.SUPP 1D), demonstrating that the model effectively denoises modality-specific signals to recover shared biological signals. Consistent with this, ISM profiles computed from MNase-seq and chemical cleavage predictions show good agreement (*r* = 0.66, Fig.SUPP 1E), indicating that the same underlying sequence features drive nucleosome positioning in both assays.

### Nucleosomes are positioned within regularly spaced islands identified by wavelet decomposition

Autocorrelation analysis of the predicted tracks yielded an NRL of *≃*187 bp, consistent with typical values observed in mammalian stem cells (Bikova et al., 2026) (Fig.SUPP1F). Genome-wide wavelet decomposition (Mallat, 1999) of the predicted MNase-seq signal revealed that nucleosomes are organized into regularly spaced positioning islands (Fig. 1B). Within *≃* 82% of the islands, a single strong ISM peak localizes within the array (Fig. SUPP 1G), suggesting that a compact sequence motif seeds the phased nucleosome pattern. Across the genome, ISM peaks tend to overlap with DNase-seq and ATAC-seq peaks (Fig. 1C), indicating that the sequence features identified by the model are enriched at accessible chromatin regions. These correlations support the view that ISM captures regulatory sequence features whose disruption alters nucleosome organization and chromatin accessibility.

### High ISM values at promoters correlate with transcriptional output

Around Transcription Start Sites (TSSs), ISM scores show a marked increase upstream of highly expressed genes (Fig. 1D). Stratification by ISM level demonstrates that promoters with stronger transcriptional output exhibit higher ISM values, consistent with regulatory sequences that both encode nucleosome depletion and promote transcription (Fig. 1D). Together, these results suggest that ISM identifies short sequence features that contribute to promoter accessibility and correlate with gene activity.

### ISM peaks reveal 5–25 bp motifs clustered within *≃*100 bp sites

To characterize the sequence elements driving nucleosome-positioning islands, we first examined the fine structure of ISM peaks. Autocorrelation analysis of the ISM signal (Fig. 2A) showed a pronounced decay within 5–25 bp. Individual ISM peaks lengths are thus consistent with TF recognition motifs, suggesting that ISM highlights discrete DNA elements whose perturbation strongly impacts local nucleosome positioning. Hereafter, we refer to these elements as NPRs. Next, we quantified the spacing between NPRs. The cumulative distribution of inter-peak distances (Fig. 2B) revealed a characteristic enrichment of NPRs within *≃*100 bp regions, demonstrating that NPRs tend to occur in compact groups rather than being uniformly dispersed along the genome. These *≃*100 bp clusters harbor multiple short regulatory motifs that collectively anchor regularly spaced nucleosome arrays.

**Figure 2.**
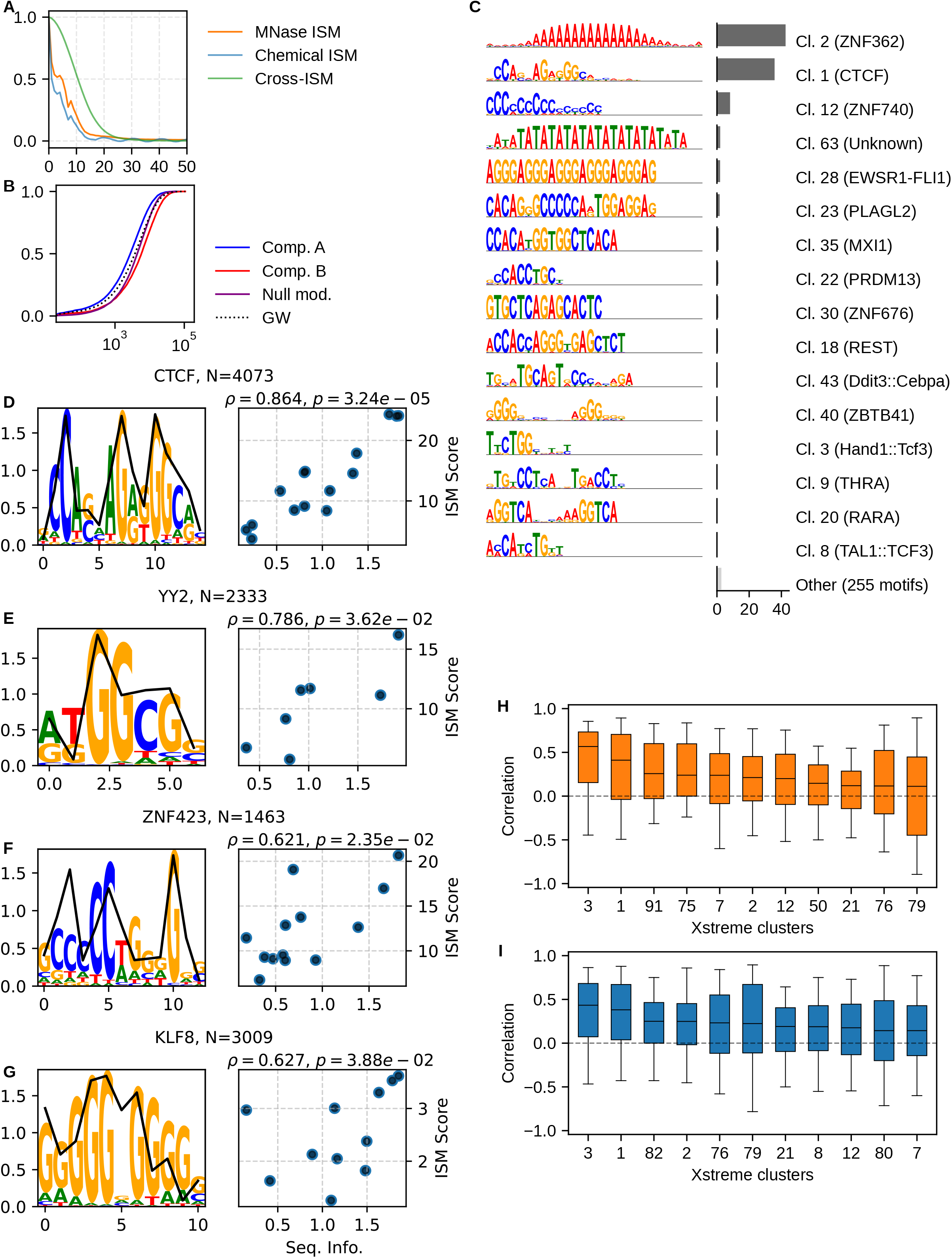
Sequence analysis of nucleosome positioning regions reveals canonical and novel motifs. (**A**) Spatial autocorrelation of *in silico* mutagenesis (ISM) scores across genomic coordinates. (**B**) Empirical cumulative distribution function (ECDF) of genomic distances between adjacent NPRs, stratified by chromatin compartment and evaluated against an exponential null model. (**C**) Representative sequence logos from the 15 most frequent clusters identified by XSTREME, annotated with their respective fraction of total NPRs. Individual clusters aggregate multiple distinct motifs (total count indicated); the top-ranking match within the JASPAR database is indicated in parentheses. (**D–G**) Sequence logos of canonical motifs derived from the JASPAR2026 database, overlaid with the mean genome-wide ISM scores computed within NPRs. The adjoined scatter plots display the relationship between sequence information content (Shannon entropy in bits; x-axis) and the mean ISM score (y-axis). (**H, I**) Box plots of Spearman’s rank correlation coefficients between position-specific information content and ISM scores computed at single-site resolution. Distributions are compared between the model trained on MNase-seq data (**H**) and the model trained on chemical cleavage data (**I**).

**Supplementary Figure 2.**
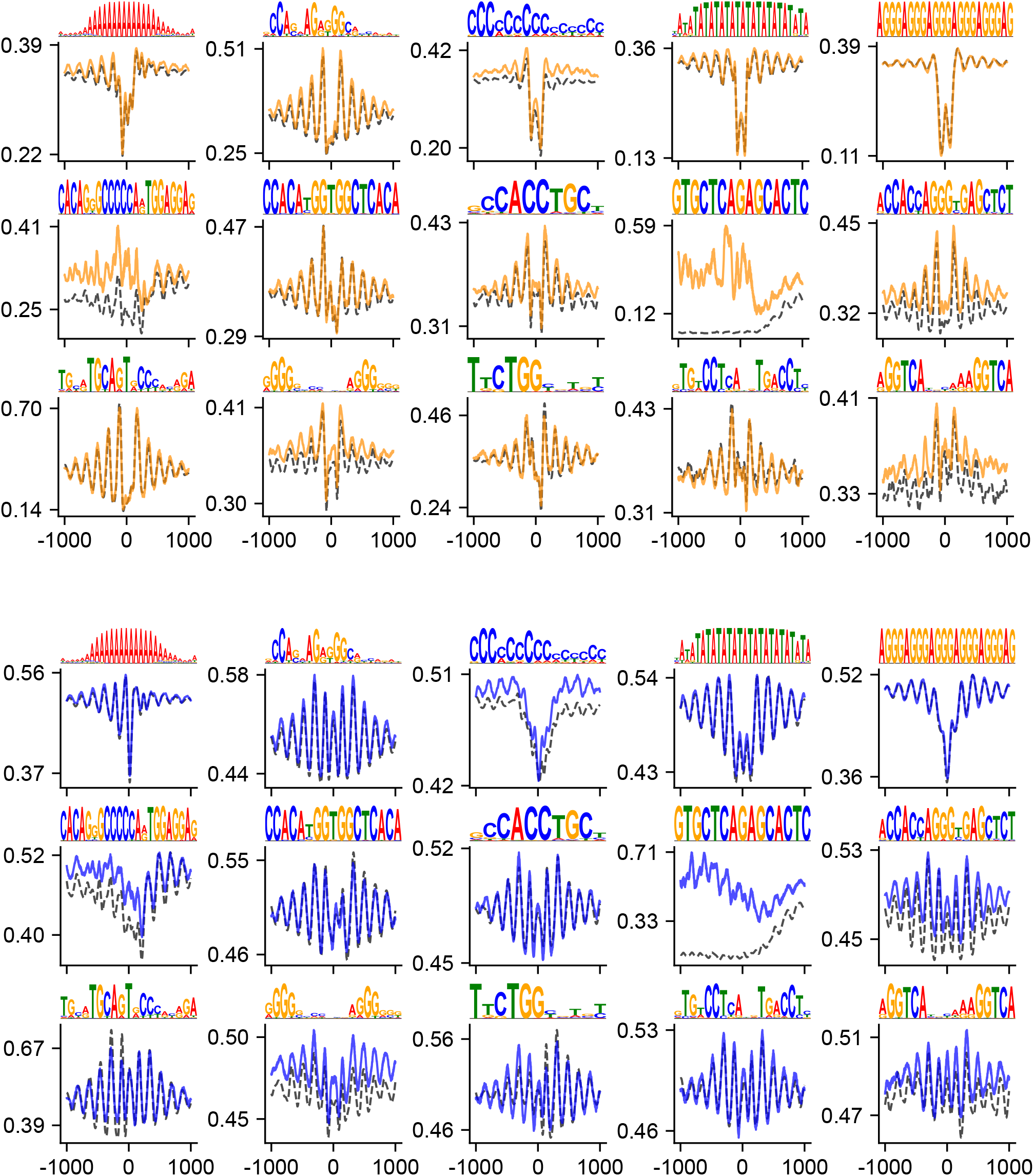
Identified sequence motifs exhibit characteristic nucleosome positioning profiles. Average nucleosome occupancy profiles computed within the flanking regions of XSTREME-identified motif clusters. Experimental MNase-seq (yellow) and chemical cleavage (blue) tracks are overlaid with their respective neural network model predictions (dashed black lines). Signal tracks are oriented strand-specifically and centered on the motif coordinates.

### NPRs are formed by both ubiquitous and cell-type-specific transcription factors

Motif enrichment analysis of NPRs identified a diverse set of TF motifs clusters contributing to nucleosome positioning (Fig. 2C). The strongest enrichment was observed for poly(A), and for CTCF, which emerged as the major organizing transcription factor of NPRs (Fig. 2C). Other high-ranking motifs included ZNF-bound sites (e.g., ZNF384, ZNF740), general regulatory factors (e.g., SP3, SPI1), and pluripotency-associated TFs (KLF2, KLF4, KLF5). This combination of constitutive architectural proteins and lineage-specific regulators indicates that NPRs integrate both universal nucleosome-organizing motifs and cell-type-restricted regulatory modules.

Consistent with this structural role, genomic regions flanking these motifs exhibit pronounced, phased nucleosome arrays , as captured by both MNase-sequencing and chemical cleavage assays and predictions (Fig. SUPP 2)

Transcription factor binding specificity is most commonly represented by Position Weight Matrices (PWMs), which quantify the contribution of each nucleotide at each position of a binding site to TF–DNA affinity. PWMs therefore provide a natural reference to interpret the base-pair–resolved sequence sensitivity captured by the ISM (Stormo, 2015). Using the motifs identified within NPRs, we computed, for each candidate TF, the correlation between the ISM signal (derived independently from MNase-seq and chemical cleavage predictions) and the corresponding PWM (Fig. 2D-G). In multiple cases, the ISM profile closely mirrors the PWM, with the strongest mutational effects aligning with the highest-information positions of the motif (Fig. 2D-G). This agreement is non-trivial because PWMs are typically inferred from TF-binding experiments (e.g., ChIP-seq or in vitro binding assays) and assume positional independence of nucleotide contributions, whereas the ISM is learned solely from nucleosome maps without explicit TF input. We summarized this analysis across TFs groups by quantifying the distribution of ISM–PWM correlations and ranking candidate TFs accordingly (Fig. 2H, I). Both MNase-based and chemical-cleavage–based ISMs yield consistently positive correlations for a substantial subset of TFs, supporting the idea that NPRs are shaped by a combination of ubiquitous factors and cell-type-specific regulators whose binding specificities define sequence features that organize nucleosomes. Because multiple TFs can recognize similar or overlapping motifs, ISM–PWM agreement provides a complementary criterion to prioritize candidate TFs when several motif matches co-occur at the same NPR: among overlapping candidates, the TF whose PWM best matches the local ISM profile is more likely to be the relevant driver of the nucleosome-positioning signal.

### DNA repeats are major contributors to NPR

Numerous repeat families from SINE, LINE, LTR, simple-repeat and low-complexity classes are enriched in NPRs (passing the 1% FDR threshold), and the strongest enrichments include both high-copy families and lower-abundance families, indicating repeat-family–specific contributions that are not explained solely by copy number (Fig. 3A). Prominent contributors include LINE elements (e.g., L1MEd, L1MEg), SINEs (e.g., B3A, B1 Mus2, B2 Mm1a), LTR retrotransposons (e.g., RLTR10, ORR1D2, ORR1E), microsatellites such as (GA)n or (T)n, and AT-rich simple repeats. Repeat classes also differ in the transcription-factor signals associated with their ISM peaks. For each repeat class, TF enrichment indicates a small number of motifs, including poly(A) and CTCF and additional TF families, accounting for most of the enrichment and a long tail of lower-frequency TF presence. The identity of these TFs changes across repeat classes (DNA, LINE, LTR, low-complexity, SINE, simple-repeat, and unique), showing that repeat-associated NPRs carry class-specific TF-associated sequence rather than a generic background of motif matches (Fig. 3B). This class-specificity is prominently driven by ZNF, which actively target repetitive templates to establish distinct structural signatures, as exemplified by their sequence-specific engagement of LTR/ERV retrotransposons (Ecco et al., 2016) and their evolutionary control over LINE lineages (Zhang et al., 2022). This provides a mechanistic rationale for the prominent ZNF-associated signals found within these specific repeat families. This supports a model in which repetitive DNA contributes to NPR formation not only by providing abundant sequence substrate, but also by embedding distinct regulatory grammars that differ between repeat classes. Consistent with this, analysis of the ISM-derived motifs within NPRs indicates that *≃* 60% of motifs resemble known TF binding sites whereas *≃* 40% remain unannotated, suggesting that repeat-associated NPRs integrate both canonical TF motifs and a substantial repertoire of currently uncharacterized short sequence signals that strongly influence nucleosome positioning.

**Figure 3.**
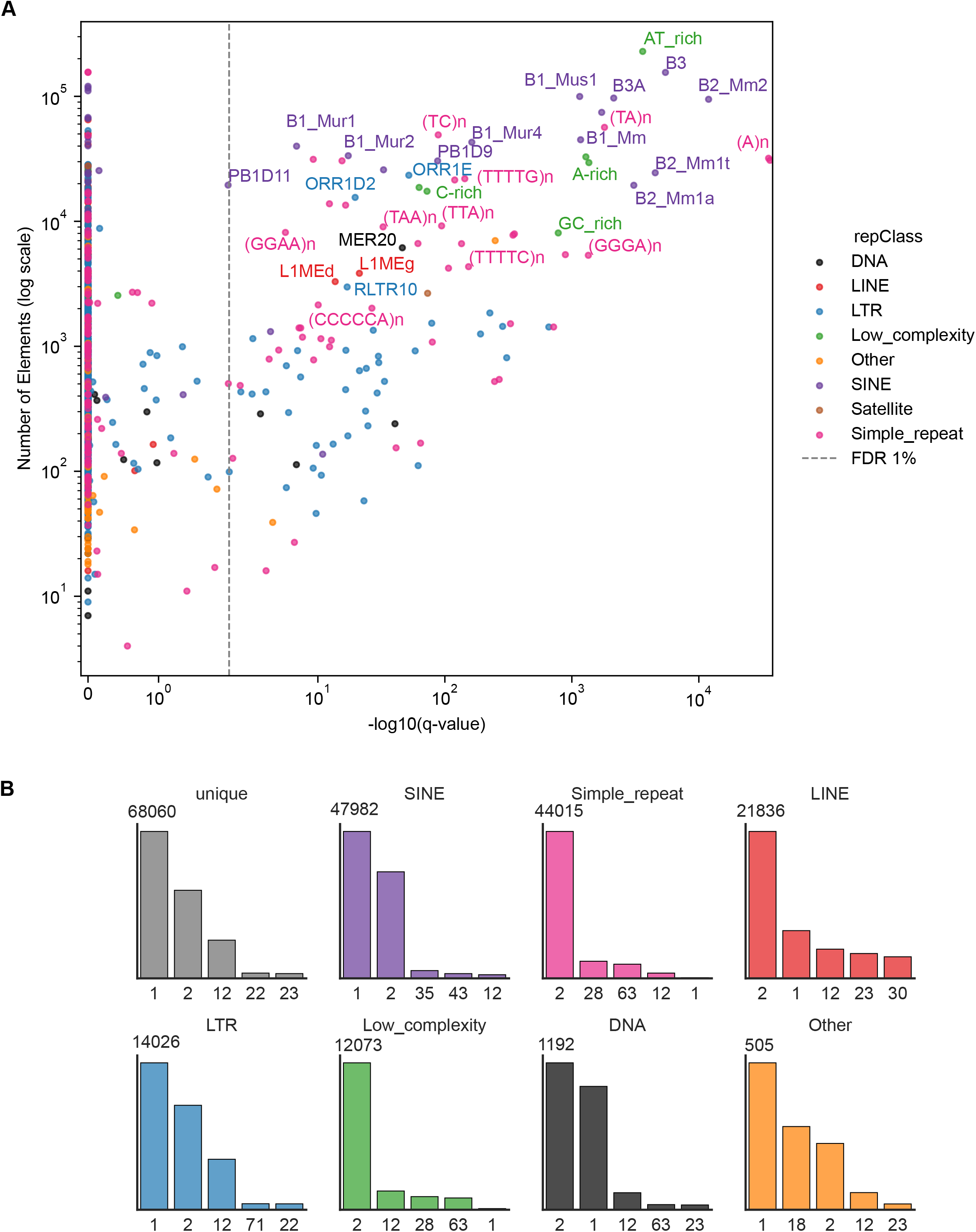
Genomic enrichment of repetitive elements within nucleosome positioning regions. (**A**) Scatter plot displaying the enrichment significance of NPRs across specific repetitive element subfamilies. The x-axis indicates the *−* log_10_-transformed *q*-value derived from Benjamini-Hochberg FDR correction, and the y-axis represents the total genomic count of elements belonging to each respective subfamily. Individual data points are color-coded according to their structural repeat class derived from RepeatMasker annotations. (**B**) Bar plot showing the absolute frequency of individual XSTREME-identified motif clusters, stratified by RepeatMasker repeat class.

Importantly, NPRs found in repetitive regions are unlikely to arise from an over-representation of repetitive elements in the training data. Since only uniquely mapped reads were retained when creating the nucleosome occupancy signals it is often equal to zero are or strongly attenuated within these regions. We therefore did not include this signal during training and all the models were trained by excluding these elements. The recovery of numerous repeat families among NPRs therefore suggests that the associated underlying sequence features associated are sufficiently informative to be learned despite their under-representation in the experimental maps.

### CTCF is a major organiser of nucleosome positioning

Approximately 35% of all NPRs correspond to CTCF motifs, consistent with its well-established role as a chromatin architectural protein that demarcates regulatory domains, positions nucleosomes, and organizes 3D chromatin loops in mammalian genome (Teif et al., 2014; Owens et al., 2019). To determine whether all CTCF motifs contribute equally to nucleosome organization, we divided high-confidence CTCF motif sites (JASPAR-score *≥* 850) sites on the test chromosome (Chr19) into two classes using ChIP-seq data: bound (*ChIP − score >* 10; n = 1232) and unbound (*ChIP − score <* 0; n = 8530). Experimental MNase-seq profiles revealed that only bound CTCF sites exhibit pronounced phased nucleosome arrays, whereas unbound motifs show no clear periodicity (Fig. 4A, red lines). This distinction is consistent with prior work demonstrating that CTCF binding can create MNase-sensitive remodeling hotspots and induce stable nucleosome positioning in embryonic stem cells Clarkson et al. (2019). Importantly, the sequence-based model faithfully reproduced the experimental phasing pattern at bound sites, while correctly predicting the absence of phasing at unbound sites (Fig. 4A, black lines). This indicates that the precise motif sequence and its local context encodes the potential to position nucleosomes. Bound CTCF motifs displayed a markedly higher ISM score for both MNase-seq– and chemical-cleavage–derived ISMs compared with unbound motifs (Fig. 4B). This result mirrors observations from Teif *et al*. showing that CTCF binding in Embryonic Stem Cells (ESCs) is largely determined by the local DNA sequence and its competitive advantage against nucleosomes (Teif et al., 2014). At the single-base level, the ISM profiles at bound sites recapitulate the CTCF PWM, with the strongest mutational effects aligned to the highest-information positions of the motif (Fig. 2D; spearman’s correlation of 0.86). De novo motif discovery using only the top ISM positions again recovered the canonical CTCF motif (Fig. 4C), reinforcing that the ISM signal directly reflects protein–DNA recognition specificity.

**Figure 4.**
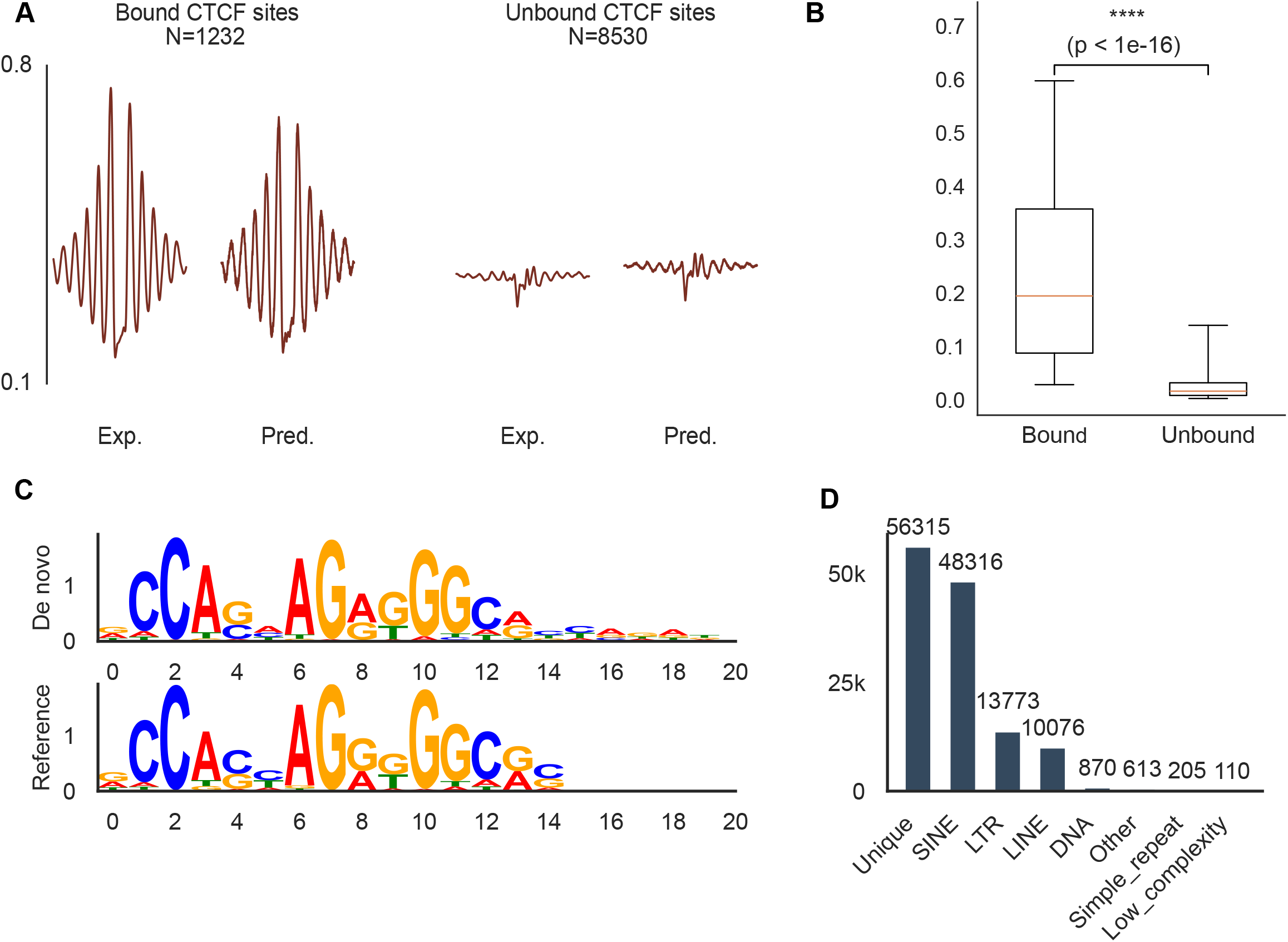
Convolutional neural network predicts functional CTCF binding sites via structural chromatin features. (**A**) Mean profiles of experimental (MNase-seq, in red) and predicted (in black) nucleosome occupancy centered on chromosome 19 CTCF binding sites (5’-3’ oriented), stratified by ChIP-seq signal (bound vs. unbound). (**B**) Box plot comparing mean ISM scores computed over bound versus unbound CTCF loci. Statistical significance was evaluated using a two-sided Mann-Whitney *U* test (*p <* 10*^−^*^16^). (**C**) Sequence logos representing the canonical CTCF motif derived from the JASPAR database alongside the *de novo* structural motif recovered by XSTREME. (**D**) Bar plot displaying the absolute distribution of functional CTCF binding sites, stratified by RepeatMasker repeat classes and non-repetitive, unique genomic sequences.

**Supplementary Figure 4.**
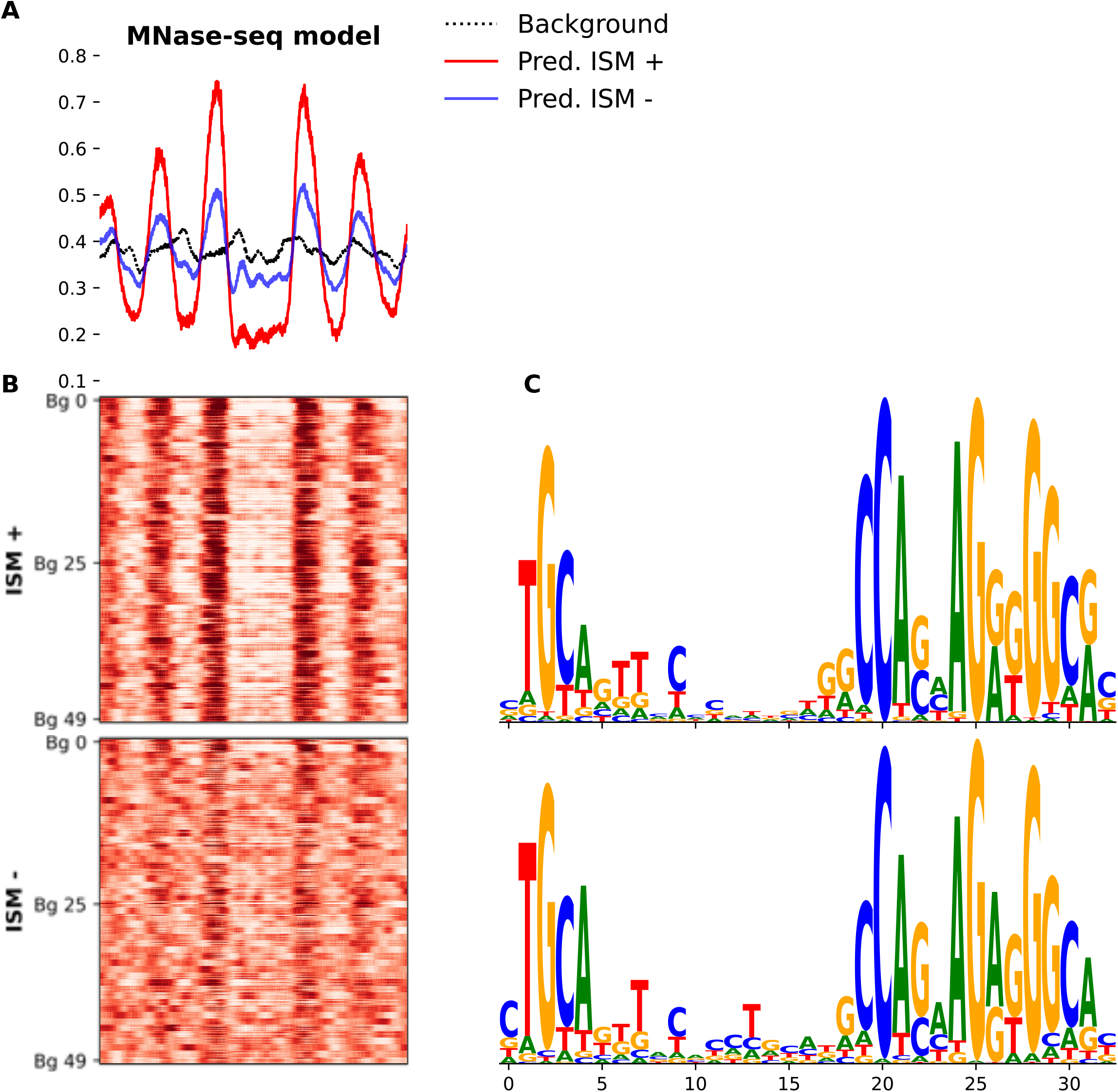
In silico motif insertion demonstrates sequence sufficiency for nucleosome positioning independently of genomic context. (**A**) Average neural network-predicted nucleosome occupancy profiles (MNase-seq) flanking genomic CTCF sites synthetically embedded into 100 distinct randomized genomic backgrounds. Loci are stratified into high-importance (ISM+, red; *n* = 320 sites) and low-importance groups (ISM-, blue; *n* = 125 sites), selected based on maximal sequence similarity to the canonical JASPAR CTCF position weight matrix on chromosome 19. The dashed black line denotes the baseline genomic background. (**B**) Corresponding heatmaps displaying predicted nucleosome occupancy profiles for individual CTCF motifs. Rows are grouped by distinct motif instances, with each instance independently evaluated across the 100 randomized backgrounds to isolate sequence-specific positioning effects. (**C**) Sequence logos illustrating the informational content (bits) of the selected CTCF binding sites for the ISM+ (top) and ISM- (bottom) groups.

To confirm this sequence-dependent effect, we predicted nucleosome occupancy across both bound and unbound motifs inserted into identical random backgrounds (see methods). Consistent with our genome-wide observations, the model specifically detects the motif sequence rather than flanking contexts: positioning CTCF motifs sustain their phasing capacity regardless of the background, whereas non-positioning variants consistently fail to establish regular arrays (Fig.SUPP 4 While bound and unbound motifs display highly similar consensus PWMs (Fig.SUPP 4C) their abilities to nucleate phased nucleosome arrays differ markedly (Fig.SUPP 4). This discrepancy indicates that standard PWMs, by assuming positional independence, fail to capture critical intra-motif dependencies. Such higher-order sequence features likely dictate the combinatorial engagement of CTCF’s 11 zinc fingers (Do et al., 2025), where subtle nucleotide co-occurrences modulate structural recognition and stable nucleosome eviction.

This is consistent with prior work arguing that CTCF binding is largely sequence-directed in ESCs (Do et al., 2025; Clarkson et al., 2019) and that differential occupancy can be modulated by chromatin state and nucleosome competition, including methylation/hydroxymethylation-dependent effects during differentiation (Soochit et al., 2021). It is also consistent with reports that CTCF shapes extended nucleosome arrays and local repeat-length changes near boundaries in ESCs, highlighting the integration of architectural proteins and nucleosome organization.

Lineage-specific expansions of CTCF binding within SINEs have been extensively documented. In mouse and rat, thousands of CTCF sites reside within B2 SINEs, each carrying strong CTCF motifs that position nucleosomes and act as insulators. Consistent with this, many of our bound CTCF motifs fall within SINE-derived CTCF-carrying repeats, while others occur in unique regions (Fig 4D).

### Microsatellites can impose sequence-dependent nucleosome positioning

Microsatellites (Short Tandem Repeats (STRs)) represent a particularly abundant class of repetitive sequences and are non-uniformly distributed along mammalian genomes, where they aggregate into dense local peaks that are frequently dominated by a single repeat motif (Xia et al., 2024). These highly variable STRs have been increasingly implicated in native regulatory roles, including modulation of chromatin structure and transcriptional output in a sequence- and length-dependent manner (Wright and Todd, 2023). Systematic annotations of transcription start sites have also revealed that these repetitive loci can possess autonomous promoter activity, serving as active platforms for pervasive non-coding transcription that can modulate nearby gene expression (Grapotte et al., 2021). Consistent with this functional potential, microsatellites are enriched within NPRs in our maps, prompting a systematic in silico assay to quantify the intrinsic nucleosome-positioning capacity of short repeat motifs. When short tandem arrays of 4-mer motifs were integrated into random backgrounds (see Methods), the average nucleosome profile showed a clear dependence on repeat number: increasing repeat copy number progressively amplified the positioning effect, indicating that repeat length acts as a gain factor for sequence-encoded nucleosome organization. Strikingly, this effect was motif-specific. For example, the homopolymeric AAAA repeat produces a strong local perturbation characterized by a pronounced depletion/valley at the repeat center together with phased oscillations in the flanking region, consistent with an imposed boundary that organizes nearby nucleosomes (Fig. 5A). In contrast, GCAA shows only a weak modulation of the average profile, indicating limited intrinsic positioning capacity under our assay conditions (Fig. 5B). Finally, GCGG produces a strong and highly periodic oscillatory pattern whose amplitude increases with repeat number, indicating robust nucleosome phasing by this motif. (Fig. 5C). Extending the analysis to all possible 4-mers revealed a broad, reproducible relationship between motif composition and positioning strength. Strong positioning 4-mers cluster at extreme GC content, such that high-positioning motifs are enriched among either very AT-rich or very GC-rich sequences, while motifs with intermediate GC content more often display weaker positioning (Fig. 5D,E). Importantly, the same qualitative trend is observed when scoring positioning using either MNase-seq–based or chemical-cleavage–based nucleosome maps, supporting that the sequence dependence is not driven solely by MNase-specific biases and instead reflects shared biological constraints on nucleosome organization.

**Figure 5.**
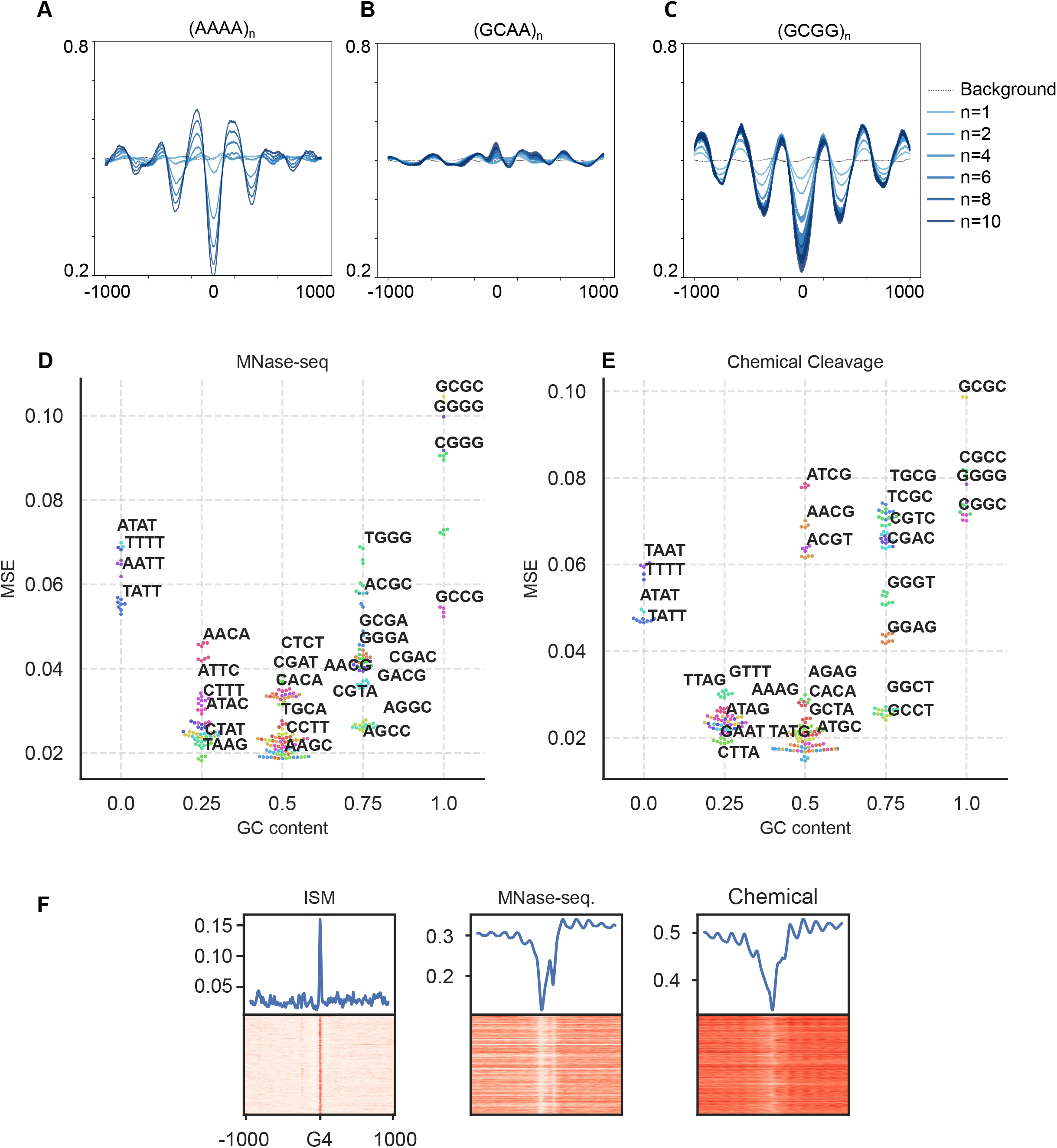
Synthetic and genomic microsatellites drive sequence-directed nucleosome positioning. (**A–C**) Metagene profiles showing the average nucleosome occupancy signal over synthetic microsatellites as a function of repeat length (blue color gradient) relative to the background (dashed black line). Representative configurations are displayed for: (**A**) (AAAA)*_n_*, (**B**) (GCAA)*_n_*, and (**C**) (GCGG)*_n_* motifs. (**D, E**) Scatter plots quantifying the impact of microsatellite sequences on nucleosome positioning, defined by the Mean Squared Error (*MSE*) relative to background tracks, and stratified by GC-content. (**F**) Heatmaps and corresponding average signal across a 2-kb window displaying ISM score alongside experimental nucleosome occupancy data (MNase-seq and chemical cleavage). Signal tracks are mapped across chromosome 19 and centered on putative G-quadruplex (G4) loci (*n* = 50182).

**Supplementary Figure 5.**
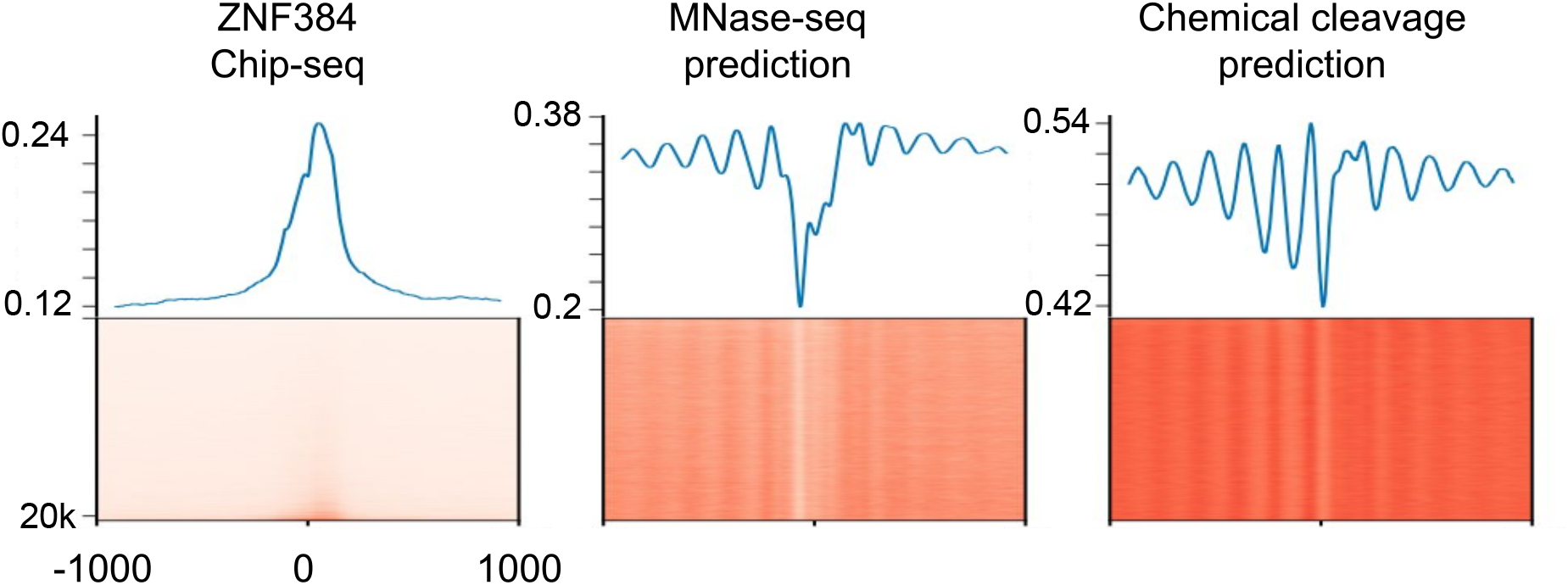
ZNF384 binding does not alter the nucleosome profile at poly-(A) sites. Heatmaps and corresponding metaplots of ZNF384 ChIP-seq signal intensity, alongside MNase-seq and chemical cleavage model predictions. Profiles are mapped across chromosome 19 and centered on homopolymeric (A)*_n_* sites associated with cluster 2.

To elucidate the biophysical basis underlying the strong positioning observed at extreme GC compositions, we first investigated whether potential G-quadruplex (G4) structures account for the positioning capacity of GC-rich microsatellites. Computational annotation of putative intra-molecular G4s revealed a significant enrichment of these non-B DNA structures within inherently strongly positioning microsatellites, which are flanked by highly ordered, periodic nucleosome arrays (Fig. 5F). Conversely, to determine whether the positioning strength of AT-rich motifs relies on trans-acting factors, we examined poly-A tracts, which frequently overlap with transcription factor binding sites such as ZNF384 (Fig.SUPP 5A). Strikingly, the characteristic nucleosome exclusion and flanking phasing profiles remained invariant upon the exclusion or depletion of ZNF384, indicating that high adenine density intrinsically dictates nucleosome positioning through DNA stiffening and steric hindrance rather than via transcription factor recruitment (Segal et al., 2006; Mavrich et al., 2008; Zhao et al., 2015).

Together, these results support a simple model in which microsatellites contribute to NPR formation through the combination of motif identity and repeat length: specific short motifs act as potent organizers of nucleosome phasing, and increasing repeat number strengthens their capacity to impose regularly spaced nucleosome arrays.

## Discussion

Understanding the sequence determinants of nucleosome positioning has been a long-standing challenge, yielding heterogeneous success across different organisms (Segal et al., 2006; Valouev et al., 2011; Teif et al., 2012). While recent deep learning architectures have achieved highly accurate predictions of nucleosome occupancy (Di Gangi et al., 2018; Amato et al., 2020; Zhou et al., 2022; Han et al., 2022), they offer limited insights into the underlying biological features they capture. Current interpretability efforts predominantly rely on analyzing *k*-mer enrichment at favored nucleosomal positions (Masoudi-Sobhanzadeh et al., 2024; Mondo et al., 2025), which fails to fully decode the regulatory logic embedded within the genomic sequence. While in yeast previous efforts were particularly productive (Routhier et al., 2021), the increased complexity of mammalian genomes has represented a bottleneck. Our work extends prior sequence–chromatin studies in three main ways: by unifying two orthogonal nucleosome-mapping assays under a single sequence model, by extracting a compact, motif-level “nucleosome positioning vocabulary” from mammalian genomes, and by showing that repetitive DNA—especially transposon families and microsatellites—provides a major substrate for that vocabulary.

First, the ability of one model to predict both MNase-seq and chemical-cleavage nucleosome maps, while the experimental tracks correlate only modestly with each other, argues that much of the disagreement between assays reflects protocol-specific noise rather than biological differences. This observation is important in mammals, where sequence contributions are often considered weaker or more context-dependent than in yeast. In yeast, reconstitution experiments and statistical models showed that intrinsic sequence preferences can explain a large fraction of in vivo nucleosome organization, providing a foundation for the “DNA-encoded” component of nucleosome positioning (Badis et al., 2008; Mavrich et al., 2008; Zhang et al., 2011). Deep learning later demonstrated that the same principle can be sharpened into base-level “mutation importance” tracks that highlight the motifs most responsible for predicted nucleosome density (Routhier et al., 2021, 2024). Here, applying a similar interpretability strategy to mESCs, we show that sequence-derived predictions from both assays correlate better with each other than the raw assays do, and that their ISM signals also align, supporting the idea that sequence features shared across datasets represent genuine nucleosome-positioning determinants rather than assay discrepancies related to intrinsic experimental biases.

Second, our ISM-based framework turns genome-wide nucleosome maps into a discovery tool for short sequence elements that nucleate phased arrays. Prior mammalian MNase-seq work emphasized heterogeneity: only a minority of loci show strong positioning, yet positioned sites can seed phased arrays whose spacing varies with cell type and chromatin state (Festuccia et al., 2019; Owens et al., 2019). Paired-end MNase-seq in humans further suggested that nucleosome arrays may be widespread (nearly half the genome in one cell type), and that many arrays reflect interplay between intrinsic sequence preferences and barrier effects (Gaffney et al. (2012); Valouev et al. (2011)). Our results refine these ideas by making the nucleation points explicit: ISM peaks reveal compact 5–25 bp features clustered within *≃* 100 *bp* NPRs, each typically associated with a single ISM peak per positioning island. This bridges “barrier” models and motif-level grammar by showing that many barriers can be localized to discrete sequence features learned directly from nucleosome maps. Importantly, NPRs correlate with DNase-seq/ATAC-seq and reflect transcriptional output around promoters, linking the sequence-defined nucleosome-organizing vocabulary to regulatory activity rather than only to physical DNA–histone energetics.

A particularly informative extension is the connection we establish between ISM signal and transcription-factor binding specificity. PWMs are the standard quantitative representation of TF sequence specificity, widely used to predict binding sites and interpret regulatory sequence. We find that the base-pair ISM profiles at NPRs often correlate with TF PWMs, and that this agreement can be used to rank candidate TFs. This is conceptually important because the model is trained only on nucleosome maps, not TF occupancy, yet it recovers TF-like specificity signatures. Practically, it offers a new way to prioritize candidate binders in regions where multiple motifs overlap or where motif databases return ambiguous TF families: among competing motifs, the PWM that best matches the local ISM profile is more likely to correspond to the factor that actually drives the nucleosome-positioning signal.

Third, our analysis places repetitive DNA at the center of mammalian nucleosome organization at the scale of positioning islands. Repeat-driven expansions of regulatory motifs have long been proposed as a mechanism for rewiring flexible mammalian gene regulation Bourque et al. (2018). CTCF provides a canonical example, with lineage-specific repeat families dispersing functional CTCF motifs across genomes (Schmidt et al., 2012). We extend this perspective from TF occupancy to nucleosome positioning by showing that diverse repeat classes (SINEs, LINEs, Long Terminal Repeats (LTRs), simple repeats and low-complexity sequences) are enriched among ISM peaks and that repeat classes carry distinct TF-enrichment signatures, implying that repeats encode class-specific regulatory grammars rather than generic motif background. The CTCF case illustrates the mechanistic logic: only motifs bound in vivo generate strong phased arrays and strong ISM signal, and the model recapitulates this bound/unbound distinction from sequence. While previous work relied on ChIP-seq data to train a model distinguishing bound from unbound states, our framework infers *in vivo* occupancy potential directly from DNA sequence and its associated nucleosome-positioning signal(Romero et al., 2024). Notably, repetitive regions contributed little to the training signal because only uniquely mapping regions were retained. Their emergence as major NPR determinants therefore argues for a genuine contribution to nucleosome organization rather than a training bias.

Finally, we provide a focused mechanistic link between microsatellite (STRs) biology and nucleosome positioning. Functional work has implicated STRs in chromatin and gene regulation in a motif- and length-dependent manner, with some repeats favoring nucleosome incorporation and others excluding nucleosomes and increasing accessibility. Our in silico repeat-length titration extends these observations by quantifying intrinsic nucleosome-positioning capacity across all 4-mers: positioning strength increases with repeat number and is motif-specific, with strong positioners enriched at extreme GC content (very AT-rich or very GC-rich), a trend observed with both MNase-based and chemical-cleavage–based scoring. This provides a simple predictive rule for how microsatellite composition and length could contribute to the formation of NPRs and to the strength of positioning islands.

In sum, our study advances the field from “sequence contributes” to “sequence elements can be enumerated, localized, and interpreted” in mammalian nucleosome organization. By integrating two nucleosome-mapping assays, extracting motif-scale NPRs via ISM, linking nucleosome sensitivity to PWM specificity, and highlighting repeat and microsatellite substrates, we provide a unified framework in which a relatively small repertoire of short DNA features—many embedded in repeats—nucleates phased nucleosome islands whose presence and strength correlate with regulatory activity.

## Methods

### Datasets

#### Nucleosome-mapping assays

##### MNase-seq data

Genome-wide nucleosome occupancy profiles in mouse embryonic stem cells (mESCs) were obtained from MNase-sequencing data generated by Festuccia et al. (2019) and are available under GEO accession GSE122589. The data were processed as in Festuccia et al. (2019) to provide paired-end fragment midpoint coordinates representing inferred nucleosome dyad positions. To construct continuous nucleosome occupancy signals, fragment midpoints were aggregated genome-wide and smoothed using a Gaussian convolution with mean *µ* = 1 and standard deviation *σ* = 15 bp. To reduce the influence of extreme values, the convolved signal was truncated at the 99th percentile and subsequently normalized to the [0, 1] range. This normalized signal was used as the MNase-seq–derived nucleosome occupancy track throughout the study.

##### Chemical cleavage data

Base-pair–resolution nucleosome positioning data derived from chemical cleavage experiments in mESCs were obtained from Voong et al. (2016) and are available under GEO accession GSE82127. These data were originally mapped to the mm9 mouse genome assembly. All coordinates were converted to the mm10 assembly using the UCSC LiftOver tool with the corresponding mm9-to-mm10 chain file, and mm10 was used consistently for all downstream analyses. Nucleosome occupancy at each genomic position was computed from nucleosome core particle (NCP) scores as described in Voong et al. (2016). Briefly, local occupancy at position *k* was calculated by summing NCP scores within a window centered on *k*, with additional center-weighting applied using a Gaussian weighting function:

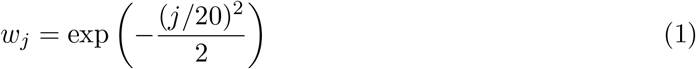

where *j* denotes the offset from the central position. This center-weighted aggregation produced a smoothed nucleosome occupancy signal that improves nucleosome boundary definition and positional robustness.

#### Reference genome

All sequence extraction and genomic analyses were performed using the mouse reference genome assembly mm10 (GRCm38). Regions overlapping undefined bases or excluded regions were omitted when applicable.

#### Transcription factor ChIP-seq data

CTCF ChIP-seq data in mESCs were obtained from GSE131356 (Owens et al., 2019) and used to classify CTCF motif instances as bound or unbound (see below). ChIP-seq data for ZNF384 in mESCs were obtained from ENCODE under accession ENCFF043AHA (https://www.encodeproject.org/).

#### RNA-seq data and transcription start sites

Strand-specific RNA-seq data in mouse embryonic stem cells were obtained from ENCODE as bigWig signal tracks for the plus and minus strands (accessions ENCFF907JEV and ENCFF373ZMQ, respectively). Genome-wide RNA-seq coverage was extracted at base-pair resolution using the mm10 reference genome.

Coordinates of transcription start sites (TSSs) were obtained from the refTSS v3.3 mouse annotation (Abugessaisa et al., 2019) mapped to the mm10 assembly. Only TSSs with complete annotation information were retained. RNA-seq values were log-transformed as log_10_(*x* + 10*^−^*^5^) for visualization and comparative analyses.

#### DNase-seq and ATAC-seq data

DNase-seq data in mESCs were obtained from ENCODE under accession ENCSR000CMW. ATAC-seq data in mESCs were obtained from ENCODE under accession GSE129673 (GSM3719303, GSM3719304).

#### Repetitive elements annotation

Genome-wide annotations of repetitive elements were obtained using RepeatMasker based on the Repbase reference library. Repeat annotations for the mouse genome (assembly mm10) were retrieved from the UCSC Genome Browser RepeatMasker table (rmsk). This dataset provides genomic coordinates and family-level classifications for major classes of repetitive elements, including LINEs, SINEs, LTR elements, and simple repeats.

For downstream analyses, repeat instances were grouped according to their annotated families and classes. When required, repeat coordinates were reoriented to match the consensus orientation of each repeat family, and their positions were expressed relative to the consensus sequence, accounting for partial or truncated elements. This standardization enabled aggregation of model predictions and sequence sensitivity profiles across heterogeneous repeat instances.

All repeat-based analyses were performed using the mm10 reference genome, and only repeat instances with unambiguous genomic coordinates were retained.

#### G-quadruplex annotation

Putative intra-molecular G-quadruplex (G4) structures within the *Mus musculus* reference genome (mm10) were computationally annotated using the g4predict pipeline (https://github.com/mparker2/g4predict). The algorithm identifies potential G4-forming sequences based on canonical folding motifs and sequence constraints, enabling systematic mapping of non-B DNA secondary structures across the murine genome for subsequent genomic integration.

### Computational Modeling and Statistical Analysis

#### Input sequence extraction and labeling

Genomic sequences were extracted from the mm10 reference genome using sliding windows of fixed length (2001 bp), centered at genomic positions sampled across the genome. Sequences were one-hot encoded over the four canonical nucleotides (A, C, G, T).

For each input sequence, the corresponding nucleosome occupancy signal was extracted and used as the prediction target. Depending on the task, targets consisted of either a single scalar value representing nucleosome occupancy at the central position or multiple values corresponding to offsets relative to the window center.

Training, validation, and test sets were defined by splitting the genome by chromosome, with chromosomes 1-17 used for training, chromosome 18 for validation, and chromosome 19 held out for testing.

Nucleosome occupancy within highly repetitive regions is frequently underrepresented or absent. We therefore excluded these regions from the training dataset, resulting in a conservative estimation of repeat-associated signals.

#### Neural network architecture

We implemented a one-dimensional convolutional neural network (CNN) using TensorFlow/Keras to predict nucleosome occupancy directly from DNA sequence. Input sequences were one-hot encoded and provided to a stack of three convolutional blocks. The first block consisted of a 1D convolution with 256 filters and a kernel size of 5, followed by max-pooling (pool size 2) and batch normalization. The second and third blocks consisted of 1D convolutions with 64 filters and kernel sizes of 11 and 21, respectively, each followed by max-pooling (pool size 2) and batch normalization. All convolutional layers used ReLU activation.

The resulting feature maps were flattened and passed through a fully connected layer with 8 units and ReLU activation, followed by batch normalization. The output layer consisted of five sigmoid-activated units corresponding to five prediction heads. These outputs represented nucleosome occupancy predictions at fixed genomic offsets of *−*500, *−*250, 0, +250, and +500 bp relative to the center of the input sequence window, thereby enabling the model to predict local nucleosome positioning profiles from a single input sequence.

Models were trained using the optimizer, loss function, and evaluation metrics described below.

#### Model training

Models were trained using the Adam optimizer with an initial learning rate of 0.001. A learning rate scheduling strategy was applied whereby the learning rate was kept constant for the first two epochs and subsequently reduced by a factor of 0.5 at each epoch.

Training used a composite loss function combining the mean absolute error (MAE) and a correlation-based term defined as 1 *− r*, where *r* denotes the Pearson correlation coefficient between predicted and observed nucleosome occupancy signals. This loss formulation encouraged both accurate signal magnitude prediction and preservation of local signal shape.

All models were implemented in TensorFlow (version 2.5) and trained on a single NVIDIA RTX 4090 GPU.

#### In silico mutagenesis and interpretability analysis

To interpret sequence features learned by the model, we performed *in silico* mutagenesis (ISM) by systematically mutating each nucleotide position within the input sequence window and measuring the resulting change in model predictions. For each genomic position, a sliding window of fixed length (2001 bp) centered on the position of interest was extracted and one-hot encoded.

At the central nucleotide of each window, the reference base was replaced by the three alternative nucleotides, generating three mutant sequences per position. Each mutated sequence, together with the corresponding non-mutated reference sequence, was passed through the trained model to obtain predictions across the five output heads corresponding to offsets of*−*500, *−*250, 0, +250, and +500 bp relative to the window center.

For each mutation, the mutational effect was quantified as the mean squared difference between the prediction for the mutated sequence and the prediction for the reference sequence, averaged across output heads. Mutational scores were computed genome-wide in a sliding-window fashion and aggregated to generate position-specific mutational sensitivity profiles.

#### Classification of bound and unbound CTCF motifs

CTCF motif instances on chromosome 19 were retrieved from the JASPAR2024 database (matrix MA0139.2) mapped to the mm10 genome assembly. Motifs were first filtered to retain high-confidence matches with a JASPAR match score *≥* 850. Each retained motif was intersected with CTCF ChIP-seq peaks using PyRanges, requiring strand-matched overlap of at least 12 bp between the motif and the nearest ChIP-seq interval; motifs with no overlapping ChIP-seq interval were retained as candidate unbound sites. Motifs were then classified as *bound* if the overlapping ChIP-seq peak had a score *≥* 10 (*n* = 1232), and as *unbound* if the ChIP-seq score was *<* 0 or no overlapping peak was found (*n* = 8530). Sites with intermediate ChIP-seq scores (0 *≤* score *<* 10; *n* = 11) were excluded from the comparison. This classification was used to stratify nucleosome occupancy profiles and ISM scores in Fig. 4.

#### Definition of nucleosome positioning regions (NPRs) and sequences analysis

Nucleosome positioning regions (NPRs) were defined from genome-wide *in silico* mutagenesis (ISM) profiles derived independently from models trained on MNase-seq and chemical cleavage data. For each genome-wide ISM track, signals were first standardized using *z*-score normalization and subsequently smoothed using a rolling mean with a window size of 12 bp to reduce high-frequency noise.

To identify regions consistently supported by both experimental modalities, the smoothed MNase-seq and chemical cleavage ISM signals were multiplied element-wise, yielding a joint ISM signal highlighting concordant sequence sensitivity. Peaks were then called on this combined signal using scipy.signal.find peaks with a prominence threshold of 1. Each detected peak was expanded to a fixed 20-bp interval centered on the peak summit (*±*10 bp), defining the final set of NPRs subjected to motif discovery using XSTREME. Discovered motifs were compared against known transcription factor binding motifs from the JASPAR database using TOMTOM. Motifs were clustered and annotated based on similarity to known transcription factor families. Enrichment analyses were performed relative to matched genomic background sequences.

#### Enrichment of NPRs in repetitive element families

To assess the association between NPRs and repetitive elements, RepeatMasker annotations (mm10; chromosomes 1–19) were intersected with NPR coordinates. Overlaps were computed using PyRanges, and for each repeat family (repName, with associated repClass and repFamily), we counted the number of unique repeat instances that intersected at least one NPR by *≥* 1 bp. This count was normalized by the total number of annotated instances for each family.

Statistical enrichment was evaluated using a right-tailed hypergeometric test under a uniform placement null model. For each repeat family, the genomic coverage fraction was computed as the union length of all annotated intervals divided by the total genome length. The genome was discretized into bins of length equal to the mean NPR size, yielding a total of *N* bins and an expected number of success bins *K* proportional to the family’s coverage fraction. Given the total number of NPRs *n* and the observed number of NPR-intersecting repeat instances *x*, significance was computed as:

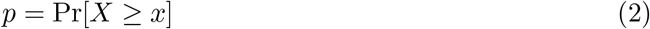

where *X ∼* Hypergeom(*N, K, n*). Multiple testing across repeat families was controlled using the Benjamini–Hochberg procedure, and enrichment was reported as *−* log_10_(*q*).

#### Continuous Wavelet Transform for Periodicity Detection

To quantify local nucleosomal periodicities from predicted genomic signal profiles at base-pair resolution, we applied a Continuous Wavelet Transform (CWT). We selected a complex Morlet mother wavelet (cmor25-0.85, bandwidth *B* = 25, center frequency *C* = 0.85). The high bandwidth parameter (*B* = 25) was chosen to maximize spectral resolution, allowing precise discrimination of nucleosomal periodic patterns from background spatial noise.

Scale boundaries were defined to cover pseudo-periods *T ∈* [150, 220] bp via *a* = *C · f_s_ · T* (with sampling frequency *f_s_*= 1 bp*^−^*^1^), spanning the characteristic range of dinucleosome and mono-nucleosomal repeat lengths. Scales *a* were linearly discretized into *N_a_* steps across this interval.

To mitigate edge artifacts prior to transformation, signal boundaries per chromosome were zero-padded. For each genomic position *b*, the localized scale-marginalized wavelet power density *P* (*b*) was computed by integrating the squared magnitude of the CWT coefficients *W* (*a, b*) across the designated scale range *A*:

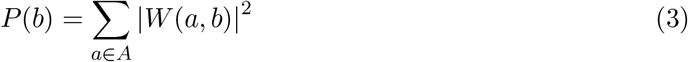

The resulting single-base energy tracks *P* (*b*) were exported as BigWig files for downstream genomic analysis.

#### Synthetic CTCF Motif Embedding and Background Perturbation

To evaluate how sequence context modulates model-predicted nucleosomal occupancy, we developed an *in silico* motif embedding framework using our MNase-seq trained model.

CTCF binding motifs (JASPAR 2024, accession MA1930.2) were extracted across the mouse genome (mm10) within strand-oriented 3000 bp windows. Motifs were stratified into footprint-positive (ISM^+^) and footprint-negative (ISM*^−^*) sets based on whether the log_10_-transformed mutation footprint signal exceeded zero within a 10 bp core window. The central 33 bp core sequences from both sets were retained for embedding.

To evaluate prediction changes across neutral sequence contexts, we generated a fixed set of 50 synthetic 3000 bp background sequences matching mouse genomic nucleotide frequencies (42% GC content). Baseline predictions for each background were derived using the MNase-seq trained model with a 2001 bp sliding window. Each 33 bp core motif (ISM^+^ and ISM*^−^*) was then individually inserted into the center of these same 50 shared background sequences, and perturbed profiles were re-predicted.

The impact of motif embedding was quantified as the integrated absolute change (*L*_1_-distance) between the baseline background prediction and the motif-inserted prediction across the 3000 bp window. Model predictions and embedded motif sets were visualized using average profile metaplots, heatmap representations across background trials, and information-content sequence logos.

#### Synthetic *k*-mer Repeat Insertion Analysis

To systematically evaluate how intrinsic DNA sequence composition and periodicity perturb model predictions independently of broad genomic context, we designed an *in silico k*-mer repeat embedding pipeline using our MNase-seq trained model.

All possible *k*-mer combinations of a given length (e.g., 4*^k^* dinucleotides) were systematically generated and tiled across multiple repetition counts (*R*) to form synthetic tandem repeat blocks. Neutral background sequences were randomly sampled from non-overlapping genomic regions to establish baseline predictions. Each *k*-mer repeat block was embedded into the geometric center of these genomic background sequences.

Model predictions were generated for both unmodified backgrounds and *k*-mer-embedded sequences using the MNase-seq trained model. The perturbation magnitude of each *k*-mer repeat configuration was calculated as the mean absolute difference (*L*_1_-distance) between the baseline background profile and the perturbed prediction across the evaluation window.

To account for structural and sequence equivalences, *k*-mers were collapsed into canonical representatives by taking the lexicographically minimal sequence among all rotational shifts and reverse complements. Perturbation scores were evaluated across varying repetition lengths, stratified by GC content, and visualized using canonical swarm plots and average profile metaplots.

## Software availability

Source code implementing the CNN models, the *in silico* mutagenesis analysis, and the NPR/motif-discovery pipeline is freely available at https://github.com/ARChE-ai/Nucleoseq under a CC-BY-SA 4.0 license.

## Data access

All newly generated data from this study — the trained convolutional neural network models (MNase-seq and chemical-cleavage), the genome-wide *in silico* mutagenesis (ISM) tracks, the predicted nucleosome occupancy tracks, and the cross-mutagenesis track — have been deposited at Zenodo under accession 22013958 (https://doi.org/10.5281/zenodo.22013958).

## Acknowledgments

M.C. was supported by a PhD fellowship from the Interdisciplinary Doctoral Program of Sorbonne Université (Interface Pour le Vivant (IPV)). This work was supported by the Agence Nationale de la Recherche (ANR) under grant no. ANR-22-CE12-0005 (*Nucleoseq* project). We thank all members of the Structure and Genome Instability Laboratory for helpful discussions.

## Author contributions

**Table 1:**
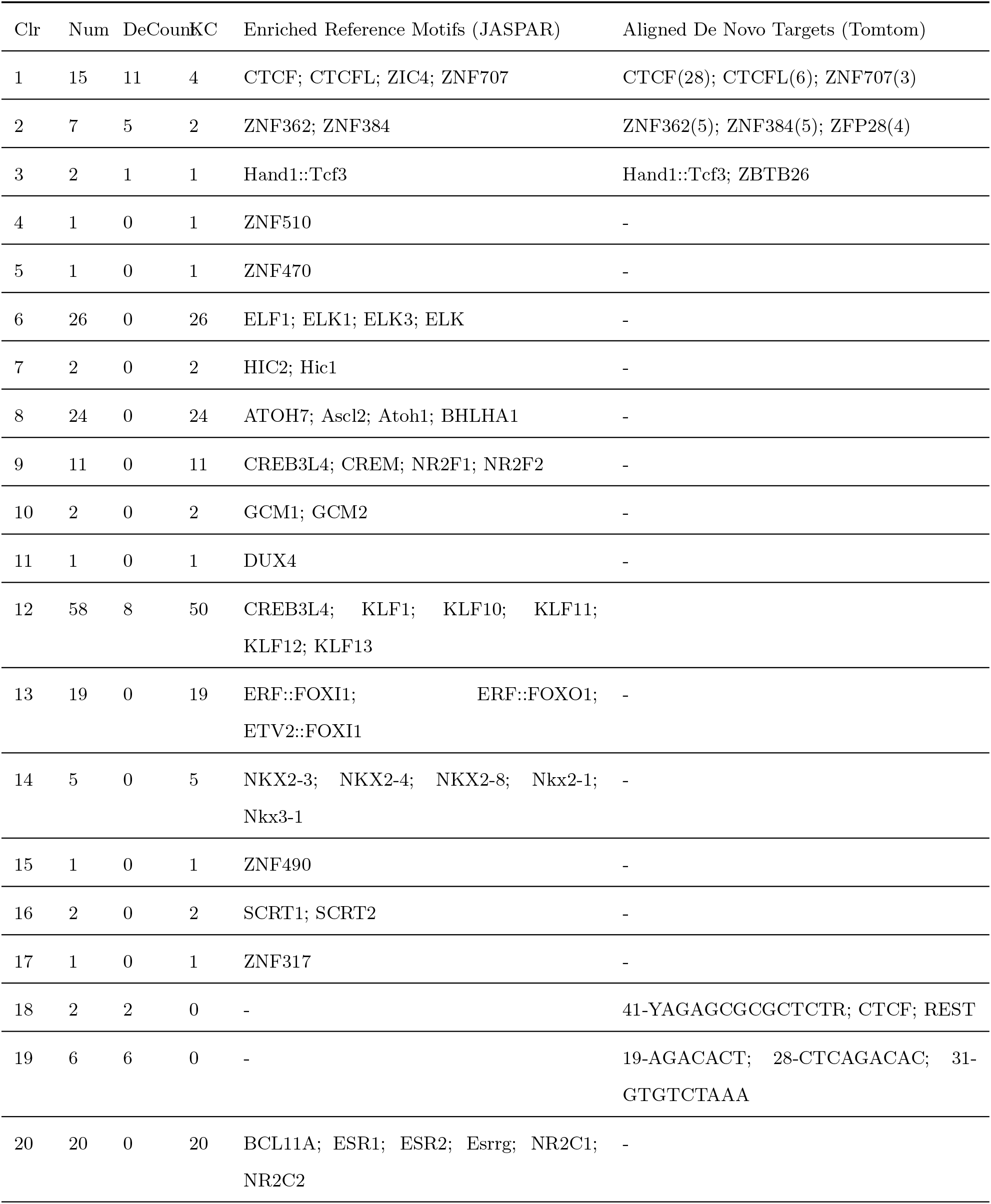
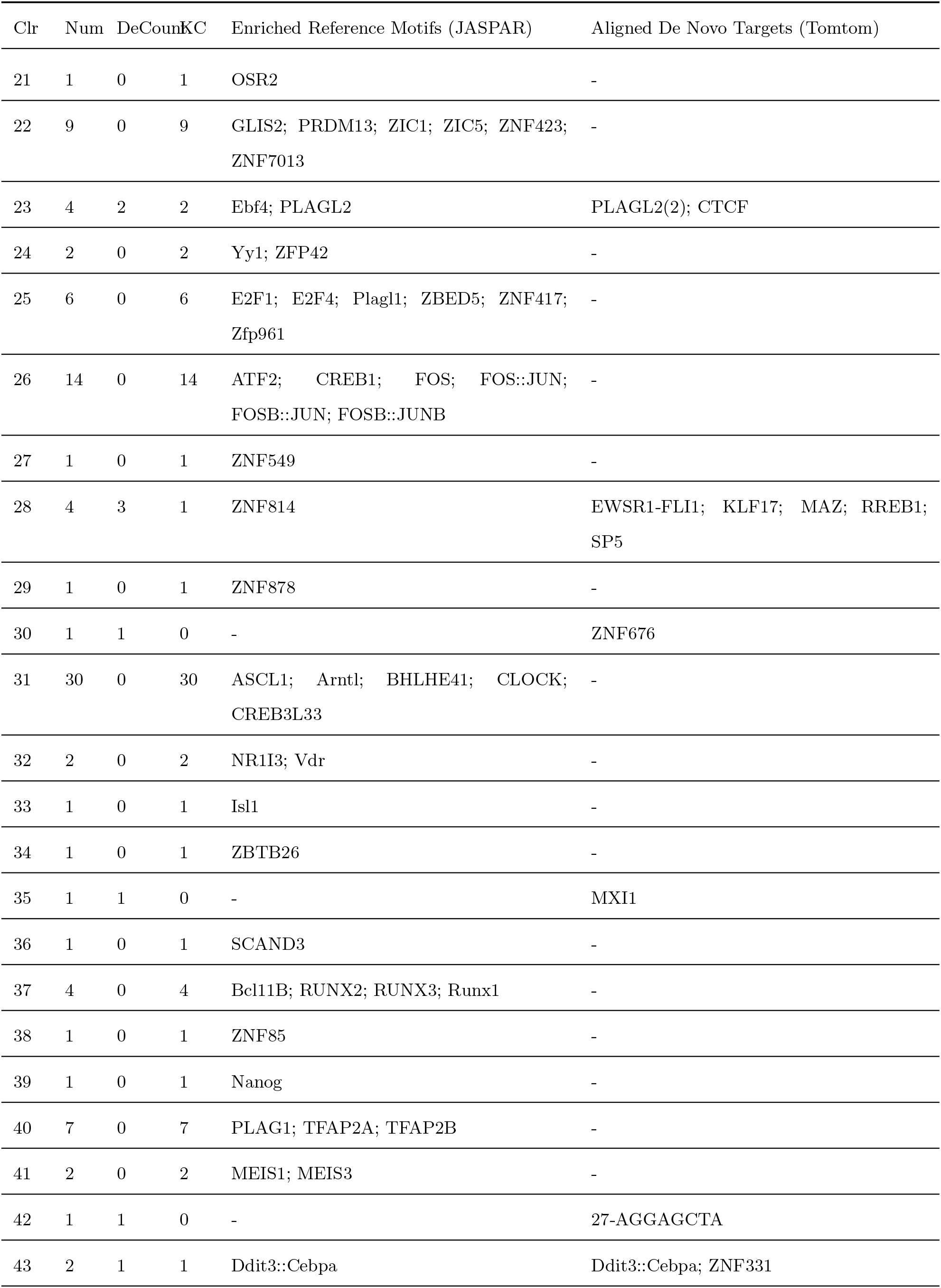
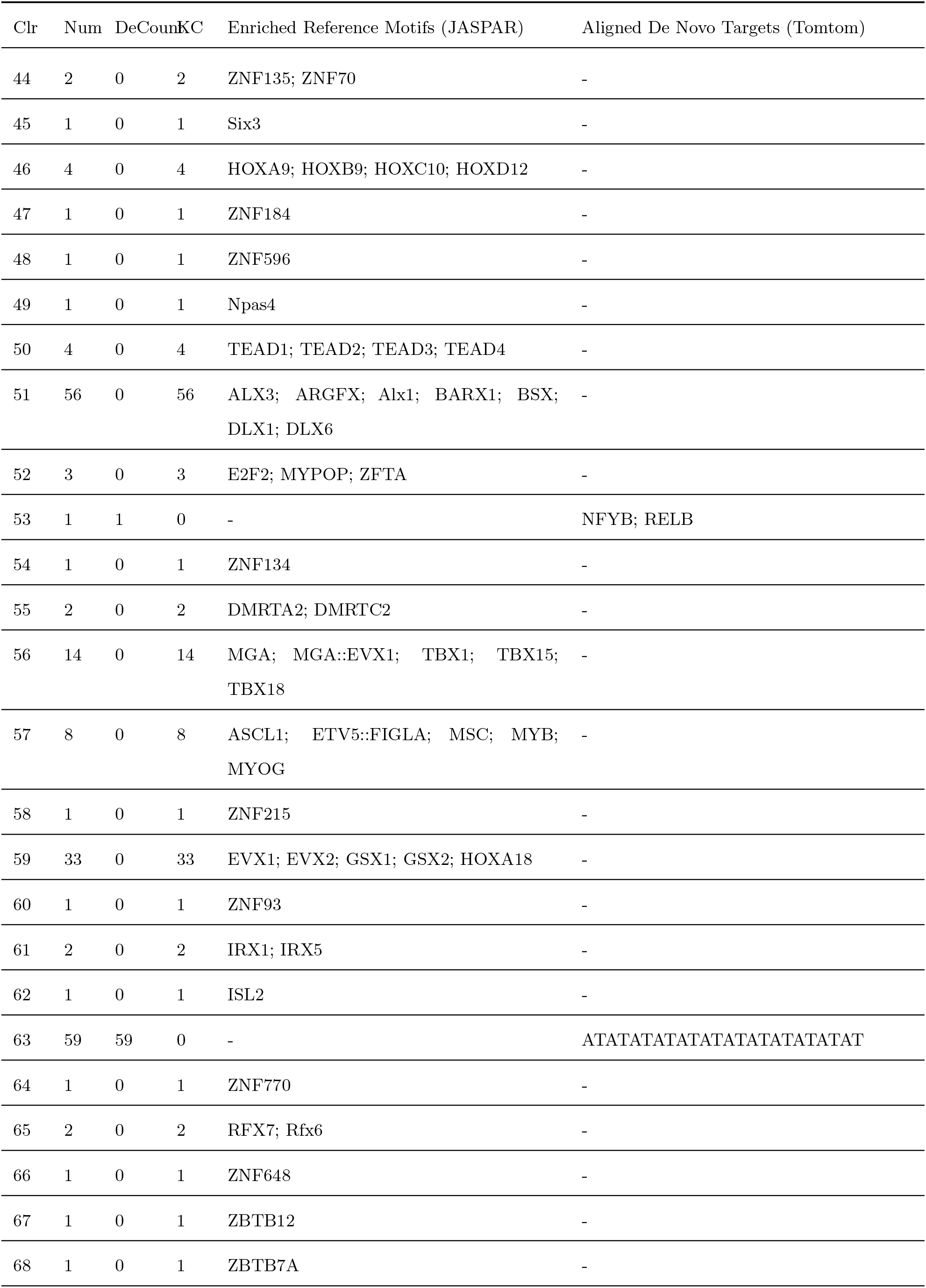
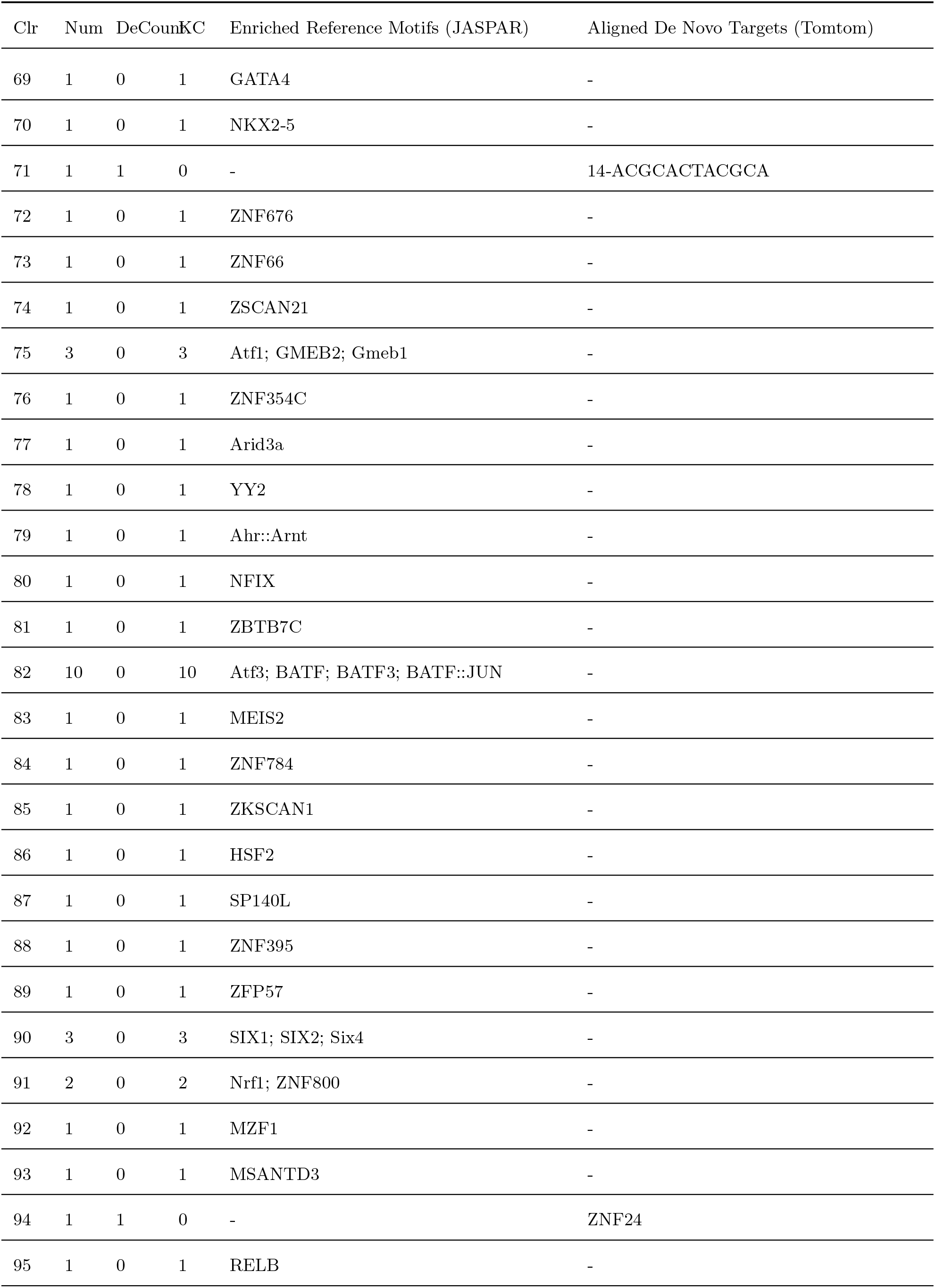
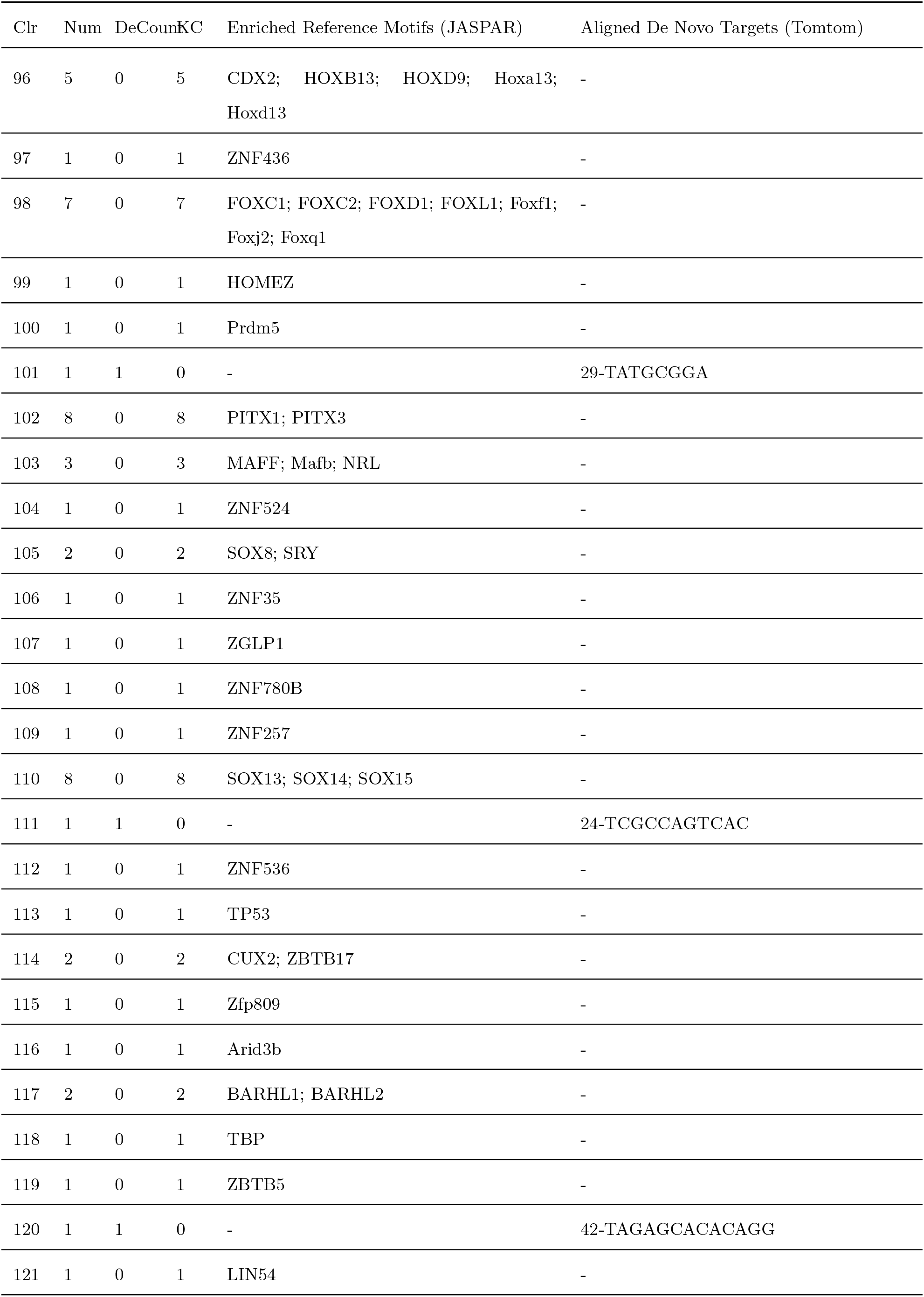
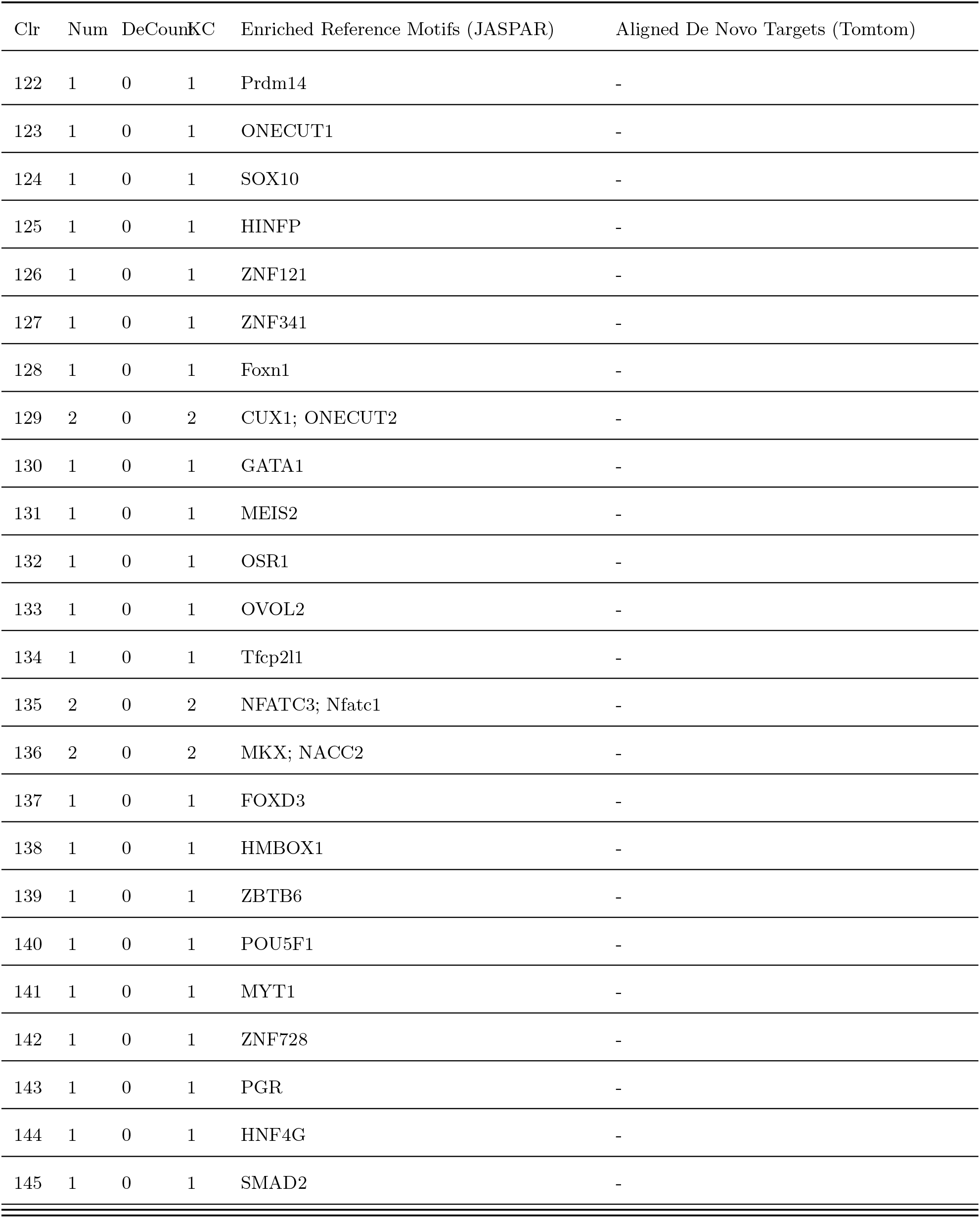
Comprehensive XSTREME cluster composition. De novo discovered matrices are listed alongside their top-ranked Tomtom alignments (unannotated sequences default to their consensus k-mer). Known enriched reference motifs are extracted directly from the JASPAR database. Absolute frequencies are indicated in parentheses; single occurrences are omitted. Elements are separated inline by semicolons.

## References

1. Abugessaisa I, Noguchi S, Hasegawa A, Kondo A, Kawaji H, Carninci P, and Kasukawa T. 2019. refTSS: A Reference Data Set for Human and Mouse Transcription Start Sites. Journal of Molecular Biology 431: 2407–2422.

2. Amato D, Bosco GL, and Rizzo R. 2020. CORENup: a combination of convolutional and recurrent deep neural networks for nucleosome positioning identification. BMC Bioinformatics 21: 326.

3. Badis G, Chan ET, van Bakel H, Pena-Castillo L, Tillo D, Tsui K, Carlson CD, Gossett AJ, Hasinoff MJ, Warren CL, et al.. 2008. A Library of Yeast Transcription Factor Motifs Reveals a Widespread Function for Rsc3 in Targeting Nucleosome Exclusion at Promoters. Molecular Cell 32: 878–887.

4. Bikova M, Clarkson CT, and Teif VB. 2026. Nucleosome spacing across cell types, diseases, and ages. Nucleic Acids Res 54: gkag074.

5. Bourque G, Burns KH, Gehring M, Gorbunova V, Seluanov A, Hammell M, Imbeault M, Izsvák Z, Levin HL, Macfarlan TS, et al.. 2018. Ten things you should know about transposable elements. Genome Biology 19: 199.

6. Chereji RV, Tolkunov D, Locke G, and Morozov AV. 2011. Statistical mechanics of nucleosome ordering by chromatin-structure-induced two-body interactions. Phys Rev E Stat Nonlin Soft Matter Phys 83: 050903.

7. Clarkson CT, Deeks EA, Samarista R, Mamayusupova H, Zhurkin VB, and Teif VB. 2019. CTCF-dependent chromatin boundaries formed by asymmetric nucleosome arrays with decreased linker length. Nucleic Acids Res 47: 11181–11196.

8. Correll SJ, Schubert MH, and Grigoryev SA. 2012. Short nucleosome repeats impose rotational modulations on chromatin fibre folding. The EMBO Journal 31: 2416–2426. Num Pages: 2426.

9. Di Gangi M, Lo Bosco G, and Rizzo R. 2018. Deep learning architectures for prediction of nucleosome positioning from sequences data. BMC Bioinformatics 19: 418.

10. Do C, Jiang G, Cova G, Katsifis CC, Narducci DN, Sakellaropoulos T, Vidal R, Lhoumaud P, Tsirigos A, Regis FFD, et al.. 2025. Binding domain mutations provide insight into CTCF’s relationship with chromatin and its contribution to gene regulation. Cell Genom 5: 100813.

11. Ecco G, Cassano M, Kauzlaric A, Duc J, Coluccio A, Offner S, Imbeault M, Rowe H, Turelli P, and Trono D. 2016. Transposable Elements and Their KRAB-ZFP Controllers Regulate Gene Expression in Adult Tissues. Developmental Cell 36: 611–623.

12. Festuccia N, Owens N, Papadopoulou T, Gonzalez I, Tachtsidi A, Vandoermel-Pournin S, Gallego E, Gutierrez N, Dubois A, Cohen-Tannoudji M, et al.. 2019. Transcription factor activity and nucleosome organization in mitosis. Genome Res. 29: 250–260. Company: Cold Spring Harbor Laboratory Press Distributor: Cold Spring Harbor Laboratory Press Institution: Cold Spring Harbor Laboratory Press Label: Cold Spring Harbor Laboratory Press.

13. Fu Y, Sinha M, Peterson CL, and Weng Z. 2008. The Insulator Binding Protein CTCF Positions 20 Nucleosomes around Its Binding Sites across the Human Genome. PLOS Genetics 4: e1000138.

14. Gaffney DJ, McVicker G, Pai AA, Fondufe-Mittendorf YN, Lewellen N, Michelini K, Widom J, Gilad Y, and Pritchard JK. 2012. Controls of Nucleosome Positioning in the Human Genome. PLOS Genetics 8: e1003036.

15. Grapotte M, Saraswat M, Bessière C, Menichelli C, Ramilowski JA, Severin J, Hayashizaki Y, Itoh M, Tagami M, Murata M, et al.. 2021. Discovery of widespread transcription initiation at microsatellites predictable by sequence-based deep neural network. Nat Commun 12: 3297.

16. Han GS, Li Q, and Li Y. 2022. Nucleosome positioning based on DNA sequence embedding and deep learning. BMC Genomics 23: 301.

17. Kaplan N, Moore IK, Fondufe-Mittendorf Y, Gossett AJ, Tillo D, Field Y, LeProust EM, Hughes TR, Lieb JD, Widom J, et al.. 2009. The DNA-encoded nucleosome organization of a eukaryotic genome. Nature 458: 362–366.

18. Kornberg RD and Lorch Y. 2020. Primary Role of the Nucleosome. Molecular Cell 79: 371–375.

19. Mallat S. 1999. A Wavelet Tour of Signal Processing. Elsevier.

20. Masoudi-Sobhanzadeh Y, Li S, Peng Y, and Panchenko A. 2024. Interpretable deep residual network uncovers nucleosome positioning and associated features. Nucleic Acids Research 52: 8734–8745.

21. Mavrich TN, Ioshikhes IP, Venters BJ, Jiang C, Tomsho LP, Qi J, Schuster SC, Albert I, and Pugh BF. 2008. A barrier nucleosome model for statistical positioning of nucleosomes throughout the yeast genome. Genome Res 18: 1073–1083.

22. Miele V, Vaillant C, d’Aubenton Carafa Y, Thermes C, and Grange T. 2008. DNA physical properties determine nucleosome occupancy from yeast to fly. Nucleic Acids Res 36: 3746–3756.

23. Mondo SJ, He G, Sharma A, Ciobanu D, Riley R, Andreopoulos WB, Lipzen A, Kuo A, LaButti K, Pangilinan J, et al.. 2025. Consecutive low-frequency shifts in A/T content denote nucleosome positions across microeukaryotes. iScience 28: 112472.

24. Owens N, Papadopoulou T, Festuccia N, Tachtsidi A, Gonzalez I, Dubois A, Vandormael-Pournin S, Nora EP, Bruneau BG, Cohen-Tannoudji M, et al.. 2019. CTCF confers local nucleosome resiliency after DNA replication and during mitosis. eLife 8: e47898.

25. Riposo J and Mozziconacci J. 2012. Nucleosome positioning and nucleosome stacking: two faces of the same coin. Mol. BioSyst. 8: 1172–1178.

26. Romero R, Menichelli C, Vroland C, Marin JM, Lèbre S, Lecellier CH, and Bréhélin L. 2024. TFscope: systematic analysis of the sequence features involved in the binding preferences of transcription factors. Genome Biology 25: 187.

27. Routh A, Sandin S, and Rhodes D. 2008. Nucleosome repeat length and linker histone stoichiometry determine chromatin fiber structure. Proceedings of the National Academy of Sciences 105: 8872–8877.

28. Routhier E, Joubert A, Westbrook A, Pierre E, Lancrey A, Cariou M, Boulé JB, and Mozziconacci J. 2024. In silico design of DNA sequences for in vivo nucleosome positioning. Nucleic Acids Res 52: 6802–6810.

29. Routhier E, Pierre E, Khodabandelou G, and Mozziconacci J. 2021. Genome-wide prediction of DNA mutation effect on nucleosome positions for yeast synthetic genomics. Genome Res. 31: 317–326. Company: Cold Spring Harbor Laboratory Press Distributor: Cold Spring Harbor Laboratory Press Institution: Cold Spring Harbor Laboratory Press Label: Cold Spring Harbor Laboratory Press.

30. Schmidt D, Schwalie P, Wilson M, Ballester B, Gonçalves A, Kutter C, Brown G, Marshall A, Flicek P, and Odom D. 2012. Waves of Retrotransposon Expansion Remodel Genome Organization and CTCF Binding in Multiple Mammalian Lineages. Cell 148: 335–348.

31. Segal E, Fondufe-Mittendorf Y, Chen L, Thåström A, Field Y, Moore IK, Wang JPZ, and Widom J. 2006. A genomic code for nucleosome positioning. Nature 442: 772–778.

32. Snyder MW, Kircher M, Hill AJ, Daza RM, and Shendure J. 2016. Cell-free DNA Comprises an In Vivo Nucleosome Footprint that Informs Its Tissues-Of-Origin. Cell 164: 57–68.

33. Soochit W, Sleutels F, Stik G, Bartkuhn M, Basu S, Hernandez SC, Merzouk S, Vidal E, Boers R, Boers J, et al.. 2021. CTCF chromatin residence time controls three-dimensional genome organization, gene expression and DNA methylation in pluripotent cells. Nat Cell Biol 23: 881–893.

34. Stormo GD. 2015. DNA Motif Databases and Their Uses. Current Protocols in Bioinformatics 51: 2.15.1–2.15.6. eprint: https://currentprotocols.onlinelibrary.wiley.com/doi/pdf/10.1002/0471250953.bi0215s51.

35. Teif VB, Beshnova DA, Vainshtein Y, Marth C, Mallm JP, Höfer T, and Rippe K. 2014. Nucleosome repositioning links DNA (de)methylation and differential CTCF binding during stem cell development. Genome Res. 24: 1285–1295. Company: Cold Spring Harbor Laboratory Press Distributor: Cold Spring Harbor Laboratory Press Institution: Cold Spring Harbor Laboratory Press Label: Cold Spring Harbor Laboratory Press.

36. Teif VB and Bohinc K. 2011. Condensed DNA: Condensing the concepts. Progress in Biophysics and Molecular Biology 105: 208–222.

37. Teif VB, Vainshtein Y, Caudron-Herger M, Mallm JP, Marth C, Höfer T, and Rippe K. 2012. Genome-wide nucleosome positioning during embryonic stem cell development. Nat Struct Mol Biol 19: 1185–1192.

38. Valouev A, Johnson SM, Boyd SD, Smith CL, Fire AZ, and Sidow A. 2011. Determinants of nucleosome organization in primary human cells. Nature 474: 516–520. Number: 7352.

39. Voong LN, Xi L, Sebeson AC, Xiong B, Wang JP, and Wang X. 2016. Insights into Nucleosome Organization in Mouse Embryonic Stem Cells through Chemical Mapping. Cell 167: 1555–1570.e15.

40. Wang JPZ and Widom J. 2005. Improved alignment of nucleosome DNA sequences using a mixture model. Nucleic Acids Res 33: 6743–6755.

41. Wright SE and Todd PK. 2023. Native functions of short tandem repeats. eLife 12: e84043.

42. Xi L, Fondufe-Mittendorf Y, Xia L, Flatow J, Widom J, and Wang JP. 2010. Predicting nucleosome positioning using a duration Hidden Markov Model. BMC Bioinformatics 11: 346.

43. Xia Y, Li D, Chen T, Pan S, Huang H, Zhang W, Liang Y, Fu Y, Peng Z, Zhang H, et al.. 2024. Microsatellite density landscapes illustrate short tandem repeats aggregation in the complete reference human genome. BMC Genomics 25: 960.

44. Zhang Y, He F, Zhang Y, Dai Q, Li Q, Nan J, Miao R, and Cheng B. 2022. Exploration of the regulatory relationship between KRAB-Zfp clusters and their target transposable elements via a gene editing strategy at the cluster specific linker-associated sequences by CRISPR-Cas9. Mobile DNA 13: 25.

45. Zhang Z, Wippo CJ, Wal M, Ward E, Korber P, and Pugh BF. 2011. A Packing Mechanism for Nucleosome Organization Reconstituted Across a Eukaryotic Genome. Science 332: 977–980.

46. Zhao H, Xing Y, Liu G, Chen P, Zhao X, Li G, and Cai L. 2015. GAA triplet-repeats cause nucleosome depletion in the human genome. Genomics 106: 88–95.

47. Zhou Y, Wu T, Jiang Y, Li Y, Li K, Quan L, and Lyu Q. 2022. DeepNup: Prediction of Nucleosome Positioning from DNA Sequences Using Deep Neural Network. Genes (Basel) 13: 1983.

